# Suspended Tissue Open Microfluidic Patterning (STOMP)

**DOI:** 10.1101/2024.10.04.616662

**Authors:** Amanda J. Haack, Lauren G. Brown, Alex J. Goldstein, Priti Mulimani, Jean Berthier, Asha R. Viswanathan, Irina Kopyeva, Jamison M. Whitten, Ariel Lin, Serena H. Nguyen, Thomas P. Leahy, Ella E. Bouker, Ruby M. Padgett, Natalie A. Mazzawi, Jodie C. Tokihiro, Ross C. Bretherton, Aaliyah Wu, Stephen J. Tapscott, Cole A. DeForest, Tracy E. Popowics, Erwin Berthier, Nathan J. Sniadecki, Ashleigh B. Theberge

**Author notes:** Corresponding authors: Nathan J. Sniadecki, & Ashleigh B. Theberge. Authors contributed equally to this work (Amanda J. Haack, Lauren G. Brown).

## Abstract

Free-standing tissue structures tethered between pillars are powerful mechanobiology tools for studying cell contraction. To model interfaces ubiquitous in natural tissues and upgrade existing single-region suspended constructs, we developed Suspended Tissue Open Microfluidic Patterning (STOMP), a method to create multi-regional suspended tissues. STOMP uses open microfluidics and capillary pinning to pattern subregions within free-standing tissues, facilitating the study of complex tissue interfaces, such as diseased-healthy boundaries (e.g., fibrotic-healthy) and tissue-type interfaces (e.g., bone-ligament). We observed altered contractile dynamics in fibrotic-healthy engineered heart tissues compared to single-region tissues and differing contractility in bone-ligament enthesis constructs compared to single-tissue periodontal ligament models. STOMP is a versatile platform – surface tension-driven patterning removes material requirements common with other patterning methods (e.g., shear-thinning, photopolymerizable) allowing tissue generation in multiple geometries with native extracellular matrices and advanced 4D materials. STOMP combines the contractile functionality of suspended tissues with precise patterning, enabling dynamic and spatially controlled studies.

## INTRODUCTION

Developing methods to generate more physiologically relevant three dimensional (3D) tissue models is important for a wide range of applications, from modeling tissue development to realizing clinical applications in tissue regeneration[1,2]. To recapitulate in vivo tissue architecture, traditional 3D cell cultures embed cells within a hydrogel consisting of natural or synthetic compositions of the extracellular matrix (ECM). However, in traditional 3D cell culture models (e.g., attached to the bottom of a well plate or in a hanging drop spheroid), it is often difficult to control mechanical cues that play a crucial role in cell morphology, alignment, proliferation, and differentiation[3,4]. To address this limitation, flexible cantilevers and pillars can serve as scaffolds for suspended 3D tissue structures that experience tensile forces[5–9]; these platforms have been used to develop engineered heart[10], musculoskeletal[11,12], and lung[13–15] tissues, as well as wound healing skin models[16]. However, suspended models typically consist of a single cell type or ECM composition, lacking the regional heterogeneity seen in natural tissues. This heterogeneity is reflected not only in the distinct junctions seen in healthy tissues (e.g., bone-ligament interfaces), but also disease processes (e.g., fibrosis) can locally alter cell types and ECM composition. Having control over the precise cellular and ECM composition within a single tissue construct increases the physiological relevance and opens new applications for modeling complex disease processes.

To achieve geometric control in 3D tissue constructs, many biofabrication techniques, such as casting[17–19], photopatterning[20,21], and 3D bioprinting[22–24], have been developed to recapitulate the complex structural and cellular composition of native tissues[25]. Of these biofabrication techniques, the casting method is widely used to generate free-standing suspended tissues (e.g., a tissue suspended between two posts in culture)[26,27]. However, the casting method is limited in geometric and cellular distribution control. Recently, 3D bioprinting has been used to generate suspended microfibers[28] via low-voltage electrospinning, and freeform structures via a sacrificial suspension bath[29–33]. These bioprinting methods require custom-engineered 3D bioprinted systems, which limit their accessibility and scalability. Microfluidic-based technologies are also widely used for their ability to introduce dynamic or perfusable flow to complex culture systems, where active flow can be used to pattern cells or cell-laden hydrogels within a microfluidic device[34–39]. It is also possible to use surface tension to drive passive flow; this type of flow has been used extensively in open microfluidic systems, eliminating the need for complex off-chip pumping systems. Open microfluidic systems are defined as having at least one channel boundary that confines the fluid flow open to the air[40–45]. Open channels can exist in multiple configurations and can be defined within solid boundaries to pattern distinct regions[40,46–49] or within fluid boundaries to pattern aqueous circuits[50]. Open-to-air channels allow for easy accessibility and manipulation of cell-ECM solutions via a simple pipetting step. Recently, open microfluidic-based 3D cell culture platforms have been realized to pattern multiple cell types or ECMs in layered 3D structures[46,51,52], for generating rail-based (i.e., a channel without walls) regional culture systems[47–49,53–56], and for patterning regions of degradable gels for temporal and spatial control of the cellular microenvironment[46,57,58]. Suspended open microfluidic systems that are devoid of a ceiling and floor have been used to generate suspended tissues composed of a uniform cell-ECM composition[51,52].

Here, we utilize suspended open microfluidics and capillary pinning to pattern 3D free-standing suspended tissues; we call this method Suspended Tissue Open Microfluidic Patterning (STOMP). STOMP interfaces with our previously described platform in which a tissue is suspended between two posts such that the tissue construct forms a bridge between these posts[26,59–62]. Previously, tissues formed using this post platform were generated with a casting method to form a tissue that is uniform in composition. In contrast, the STOMP platform uses a removable 3D-printed patterning device containing an open microfluidic channel that interfaces around the two posts. This method offers the unique advantage of spatially controlled material and cell composition by employing capillary pinning features along the open channel. Using surface tension to drive flow and capillary pinning to stop flow, we can achieve control over the placement of tissue components to pattern multiple regions within a single suspended tissue. After patterning, we either take advantage of cell compaction allowing for the tissue to pull away from the patterning walls, or employ a degradable poly(ethylene glycol) (PEG)- based hydrogel patterned in an open microfluidic channel within the STOMP channel walls to dissolve the channel wall, allowing for gentle removal of the STOMP device and revealing a suspended tissue. This system provides the capability to model complex biological features, such as border regions between tissues (e.g., disease and healthy tissue borders) and tissue junctions (e.g., bone-ligament interfaces).

## RESULTS

### STOMP is a versatile tool for achieving geometric, volumetric, and compositional control over a diverse set of materials and tissues

Free-standing engineered tissues suspended between two posts are commonly generated using a casting technique where posts are placed upside down within a well plate that contains a mold where a cell - laden hydrogel precursor solution is pipetted[62]. STOMP adds upon these existing techniques, enabling spatial control over tissue composition. To achieve this control, STOMP uses a pinning technique, where a geometric discontinuity acts as a pinning feature that opposes the driving capillary forces that advance the fluid front, thereby immobilizing the capillary flow at a predefined location[63–67] (Figure 1). Further, the use of an open channel (i.e., a channel without a ceiling and floor) allows for access so different combinations of cell-hydrogel precursors can be pipetted into different regions. The fluid fronts on either side of a pair of pinning features come together to form a contiguous tissue with distinctly different regions. The pinning features of STOMP can be placed anywhere along the length of the open channel. Moreover, multiple pinning features can be placed in one tissue, generating more than two regions. To illustrate the versatility of possible geometries, we show patterning of one, two, and three regions with colored agarose (Figure 1).

**Figure 1.**
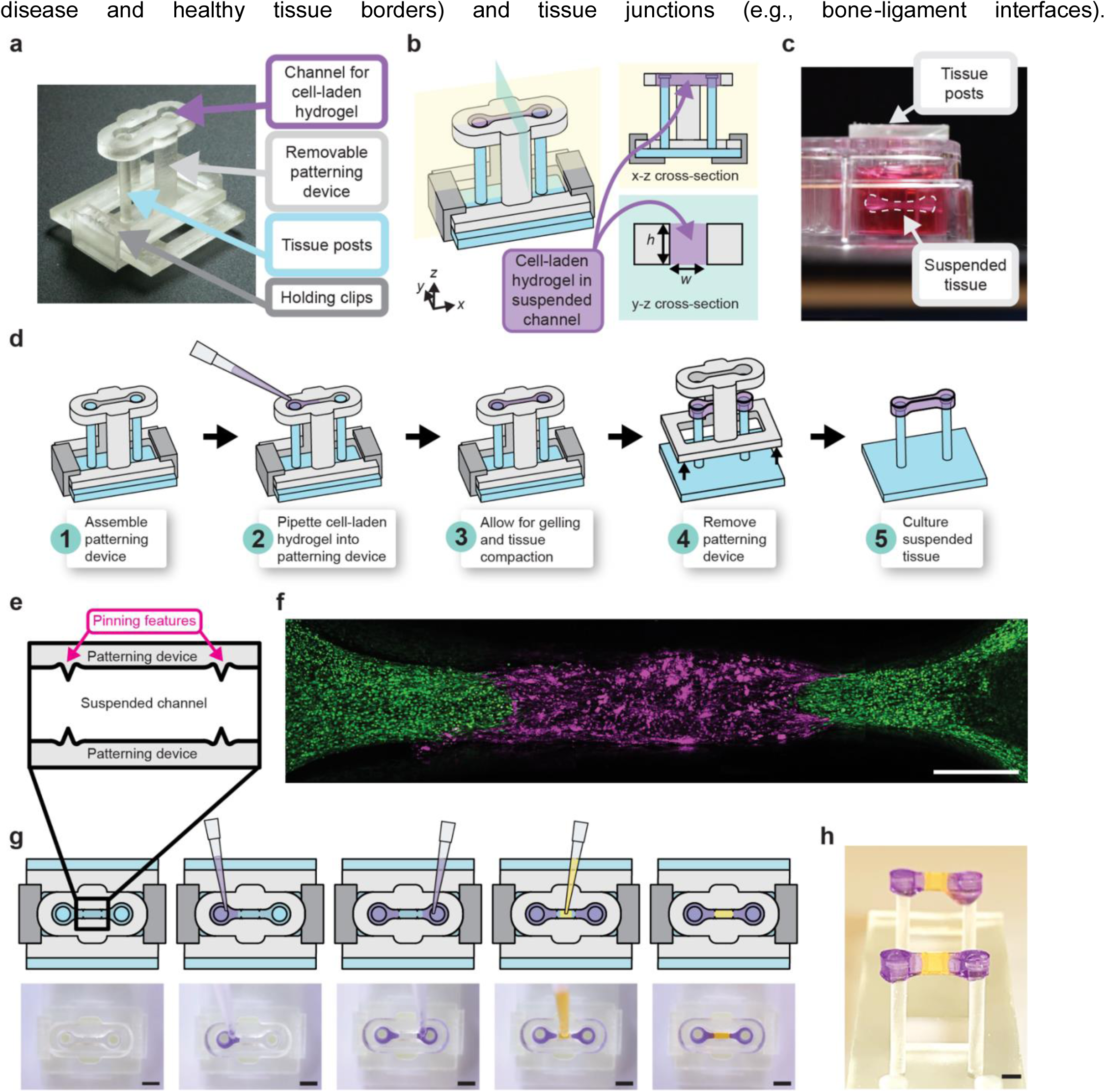
Workflow of generating single and multi-region suspended tissues using the STOMP platform. a) Image of the STOMP platform, which includes a removable patterning device containing an open channel that interfaces with a pair of vertical posts. The patterning device is held in place to the base for the posts with holding clips. b) Schematic of the STOMP platform. A cell-laden hydrogel is pipetted into the open channel, where it flows via surface-tension driven forces across the open channel and anchors onto the suspended posts, thus generating a free-standing suspended tissue. c) Side view of the resulting suspended tissue cultured in a 24-well plate. d) Workflow of patterning a tissue composed of a single region, where the composition is the same across the tissue. e) Top-down view of the capillary pinning features along the open channel that are used to pin the fluid front. f) Fluorescent image of patterned 3T3 mouse fibroblast cells laden in a fibrin hydrogel using STOMP. The outer region of 3T3 cells were dyed by CellTracker Green (green) and were pipetted first. The middle region of 3T3 cells were dyed by CellTracker Red (magenta). Scale bar is 500 μm. g) Workflow of patterning tissues comprising three distinct regions. Corresponding video stills show patterning of purple-colored agarose in the outer regions first, followed by patterning yellow-colored agarose in the middle region. Full video can be seen in Supplementary Video 1. All scale bars are 2 mm. h) Side view image of multi-region agarose suspended hydrogel construct. Scale bar is 2 mm

STOMP uses three components: (1) two vertical posts that suspend the final tissue as a bridge between them, (2) a removable patterning device containing an open microfluidic suspended channel designed to surround the two posts, and (3) holding clips to secure the assembly together (Figure 1a-b). Once assembled, we pipette a cell-laden hydrogel precursor into the open channel (Figure 1d). The solution flows throughout the open channel via spontaneous capillary flow. Once the cell-laden hydrogel solidifies and the traction forces of the embedded cells cause the construct to pull away from the walls of the patterning device, we remove the patterning device and the construct is cultured as a free-standing suspended tissue.

The precursor solution is held suspended between the walls of the open channel of the patterning device via surface tension as the solution transitions from liquid to gel state[40]. After gelation, the entire assembled patterning device is placed in a 24-well plate with media (Figure 1c). To prevent the tissue from sticking to the walls, we incubate the channel with 1% bovine serum albumin (BSA) prior to pipetting the precursor solution. T-shaped caps on each of the posts anchor the tissue such that it stays suspended between the tips of the posts. After visible compaction, the patterning device is gently lifted, thereby releasing the tissue while it stays suspended between the two posts. In this work, we have generated suspended tissues with STOMP composed of multiple hydrogel and ECM types including type I collagen, fibrin, matrigel, agarose and custom-engineered enzymatically degradable poly(ethylene glycol) (PEG). We have also used multiple cell types including fibroblasts, periodontal ligament cells, and cardiomyocytes. We demonstrate using two sets of pinning features (Figure 1e) with 3T3 mouse fibroblast cells dyed by CellTracker Green CMFDA and CellTracker Red CMTPX patterned to make a three-region configuration (Figure 1f). This workflow requires three pipetting steps, where different precursor solutions on either side of the pinning feature will meet (Figure 1g). We further demonstrate this spatial patterning of three separate regions in a free-standing tissue with colored agarose (Figure 1h).

By adding this regional control we enable in vitro models to investigate heterogeneity within a single tissue (e.g., diseased adjacent to healthy tissue), or even the investigation of multiple tissue interfaces within a single construct (e.g., bone adjacent to ligament tissue). STOMP also offers volumetric control in tissue size. In open microfluidic systems, surface tension is the main driving force of flow[40–43,68]. By changing the dimensions of the patterning channel, we can change the shape of the generated tissue as long as the cross-sectional dimensions of the channel follow the equation necessary for spontaneous capillary flow (Eq. 1), where pf is the free perimeter and pw is the wetted perimeter of the cross section[40]. Equation (1) can be simplified to equation (2) for a suspended channel geometry where w is the width (free perimeter) of the channel, h is the height (wetted perimeter) of the channel, and θ is the contact angle of the hydrogel on the channel surface[40,46]. These dimensions can be visualized in the x-z cross-section in Figure 1b.

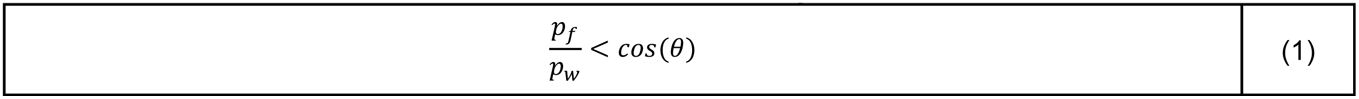

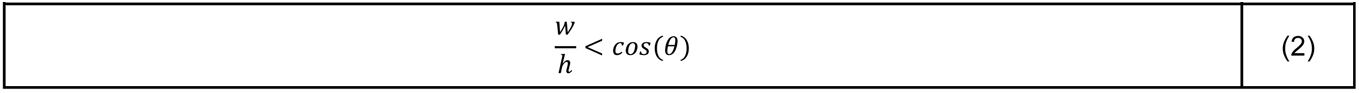

With STOMP, dimensions smaller than the capillary length of a fluid – where surface tension dominates over gravity – are preferred to maintain a uniform cross-section of the hydrogel fluid front along the channel. At larger geometrical dimensions, droplets can form within the channel, thereby preventing the complete filling of the open channel (i.e., wetting of the hydrogel along channel walls). Therefore, by maintaining the conditions for spontaneous capillary flow in a suspended channel (Eq. 2), we scaled our current STOMP design to achieve tissue volumes between 30-100 μL of hydrogel precursor, depending on channel dimensions. With customizable channel geometries, STOMP can be used to generate tissues of various sizes. For instance, a smaller volume may be advantageous when using precious cells or designer hydrogel material, and a larger volume may be advantageous for longer culture periods in tissues that compact and remodel their matrix.

### Theoretical modeling of capillary pinning features for control over 3D geometry within a suspended tissue construct

To pattern distinct regions, we use pinning features to stop the fluid front, thus creating a defined border region. However, when a second solution is pipetted into the channel on the other side of the pinning feature, the pinning must intentionally be broken, allowing the two fluid fronts to meet to form a continuous tissue. Here, we analyze two pinning feature designs we refer to as convex “vampire” or concave “cavity” pins. The convex “vampire” pinning feature design comprises opposing reliefs resembling “teeth”, with the reliefs extending along the height of the channel (Figure 2a). We note that α is the angle formed by the triangular shaped relief, θ is the contact angle of the hydrogel with the channel wall, and w_1 is the distance between the two opposing reliefs of the convex “vampire” pins (Figure 2a). If γ denotes the surface tension of the liquid, the Laplace pressure, P, of the liquid is given below (Eq. 3), where r is the curvature radius of the pinned interface.

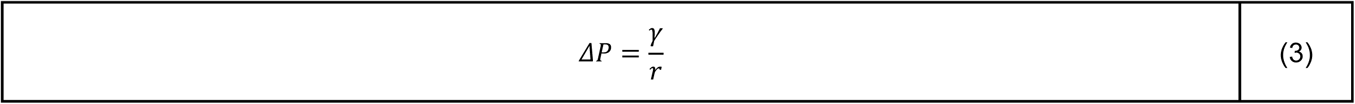

**Figure 2.**
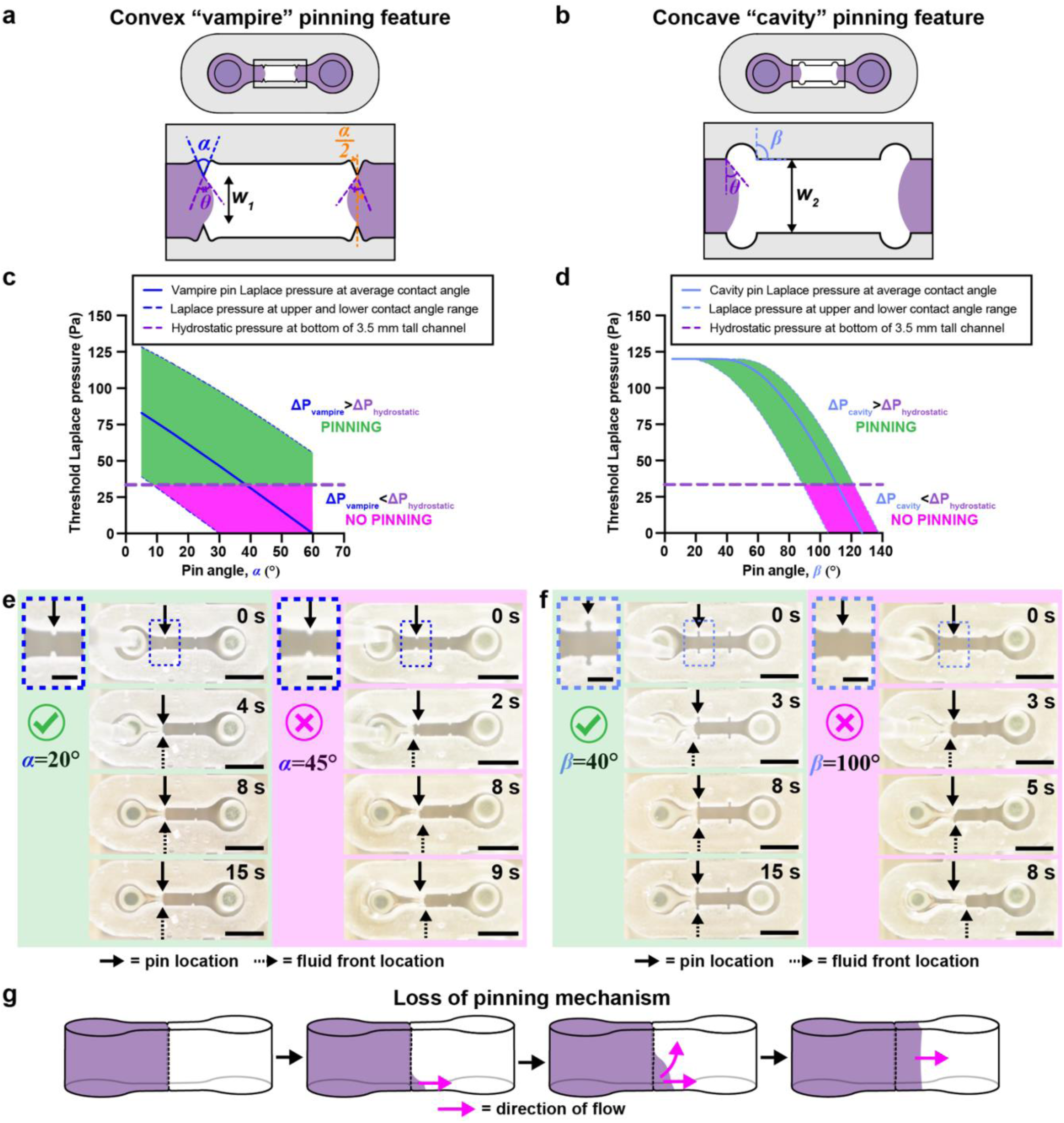
Characterization of capillary pinning features used in STOMP to generate multi-region suspended tissues. Geometric considerations of a) vampire and b) cavity pinning feature designs. Graphs of maximum Laplace pinning pressure plotted against c) *α,* pin angle of the vampire feature and d) *β*, pin angle of the cavity feature. Solid lines represent a Laplace pinning pressure (ΔP_vampire_ or ΔP_cavity_) for a contact angle of *Ө =* 30°, which is the average contact angle for 5 mg/mL collagen on 3D-printed resin treated with 1% BSA at room temperature for one hour. Upper and lower bounds of the Laplace pinning pressure is calculated based on the largest measured contact angle (*Ө =* 48°) and the smallest measured contact angle (*Ө =* 15°) on 1% BSA-treated 3D-printed resin. If the Laplace pinning pressure is greater than that of the hydrostatic pressure (ΔP_hydrostatic_ = 33.5 Pa) then pinning is predicted to occur (shaded in green). If the Laplace pinning pressure is less than ΔP_hydrostatic_, then pinning will not occur (shaded in magenta). e) Representative video still images of 23 μL of a 5 mg/mL precursor collagen solution pipetted into two different 1% BSA treated STOMP devices containing the vampire pinning features; left four images use devices with *α* = 20° and right four images use devices *α* = 45°. f) Representative video still images of 23 μL of a 5 mg/mL precursor collagen solution pipetted into two different 1% BSA treated STOMP devices containing the cavity pinning features; left four images uses devices with *β* = 40° and right four images use devices with *β* = 100°. Scale bars on insets are 1 mm. All other scale bars are 3 mm. g) Visualization for the loss of pinning mechanism, where a pink arrow indicates the direction of flow of the purple hydrogel, with pinning first lost at the bottom of the channel.

At the pinning limit (i.e., the maximum interface the fluid bulges before pinning is broken), the interface forms the angle θ with the triangular edge[64–67,69] (Figure 2a). At this pinning limit, the relationship between the geometry of the convex “vampire” pins (Figure 2a) and the contact angle θ is described by equation (4); a more detailed derivation is in the Supplementary Text section 1, and geometric considerations are illustrated in Supplementary Figure 4.

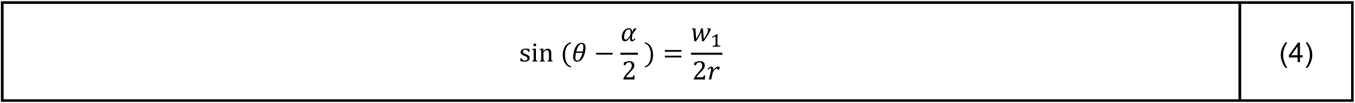

Substituting equation (4) into equation (3) yields the threshold Laplace pinning pressure, above which pinning is lost in the convex “vampire” pinning feature design (Eq. 5).

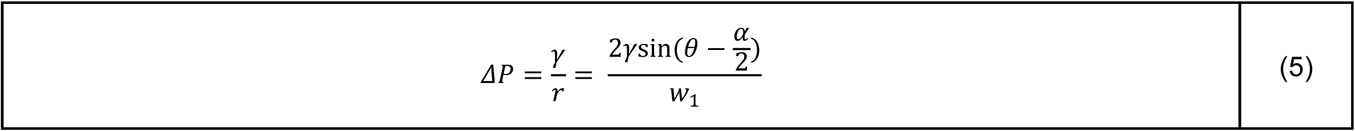

The concave “cavity” pinning feature comprises two facing cuts into the open channel resembling circular cavities, with the cut extending along the height of the channel (Figure 2b). We note that *β* is the angle formed by the line tangent to the circular cavity and the wall of the channel, θ is the contact angle of the hydrogel with the channel wall, and w2 is the width of the open channel (Figure 2b). Equation (6) describes the geometric relations of the concave “cavity” pinning feature and the contact angle at the pinning limit. A more detailed derivation is in the Supplementary Text section 1 and geometric considerations are illustrated in Supplementary Figs. 5-6.

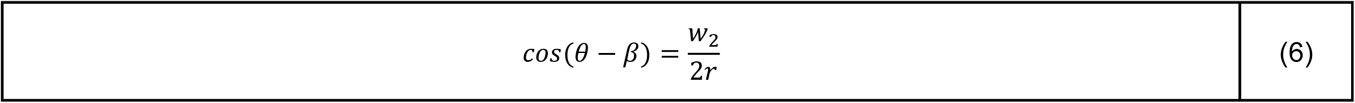

Substituting equation (6) into equation (3) yields the threshold Laplace pinning pressure, above which pinning is lost in the concave “cavity” pinning feature design (Eq. 7).

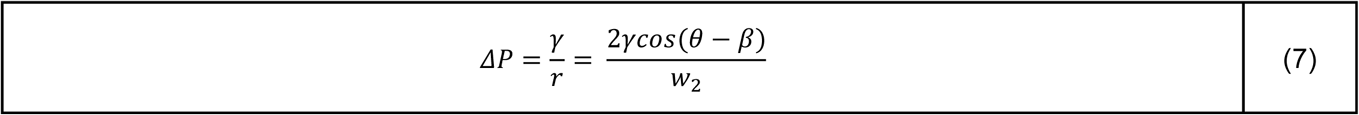

Because gravity has a small, but non-negligible effect, we must also consider hydrostatic pressure at the pinning interface. For both pinning feature designs, hydrostatic pressure at the bottom of the channel is given below in equation (8), where *ρ* is the density of the hydrogel, *g* is the gravitational constant, and *h* is the height of the channel. The effect of gravity on our system is further described in the Supplementary Text section 2 and Supplementary Fig. 7.

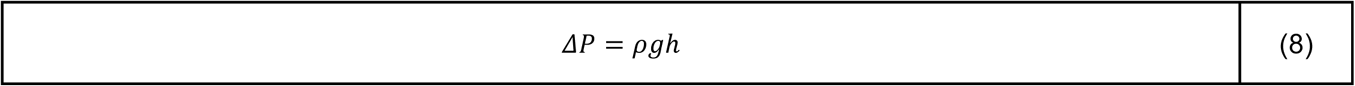

The pinning is determined by the fluid pressure at the pinning site. Pinning remains stable if the fluid pressure is lower than the threshold Laplace pinning pressure. The highest pressure occurs at the bottom of the channel where hydrostatic pressure is maximum. Therefore, the stability of pinning is dictated by the conditions at the bottom of the device. If pinning is lost at the bottom, it leads to a gradual loss of pinning along the entire vertical ridge. Figure 2g depicts a representative illustration of what the fluid front looks when pinning of the first pipetted region fails.

According to equations (5) and (7), the angles α and *β* of the convex “vampire” and concave “cavity” pins, respectively, affect whether pinning is maintained or lost (Figure 2a-b). We plotted a range of these angles to generate a theoretical prediction of the threshold Laplace pinning pressure using the parameters for 5 mg/mL collagen, with a 1% BSA treated 3D printed resin surface (Figure 2c-d). The average contact angle was found to be 30°±2° (n=20). We also considered how the range of the contact angle (15-48°) affects the theoretical modeling (see Supplementary Table 1). Using the values of the different parameters in Supplementary Table 3, we plot the threshold Laplace pinning pressure for different pinning angles (α and *β*) for the convex “vampire” (Figure 2c) and concave “cavity” design (Figure 2d), respectively.

Considering the average contact angle found for type I collagen on 1% BSA treated resin (30°), the pinning angle above which pinning is lost is when α is 35° for the convex “vampire” pinning feature; this is when Eq (5) is equal to Eq (8). For the concave “cavity” pinning features, pinning is lost when the pinning angle *β* is between 100° and 105°; this is when Eq (7) is equal to Eq (8). These angles are where the Laplace pinning threshold line crosses the hydrostatic pressure line (Figure 2c-d). When the maximum hydrostatic pressure is greater than the threshold Laplace pinning, pinning in the first pipetted region will likely fail. Loss of pinning will occur if the pressure at the fluid front exceeds the threshold Laplace pinning. The hydrostatic pressure at the bottom of the device is constant and does not depend on the geometry of the pinning features. Therefore, when the threshold Laplace pinning curve crosses below the hydrostatic pressure line, pinning will fail because the total pressure, which is dictated by the hydrostatic pressure, exceeds the threshold Laplace pressure.

When tested experimentally, we found that the success of pinning reflects these theoretical predictions while also demonstrating the variability in the system, i.e., success varies due to the large range of possible contact angles (Supplementary Figure 2). When measuring the contact angle, we found variability between separate parts, with a relatively large range of contact angles (from 15° to 48°). This large range of the contact angles are due to multiple factors, including the variability of the BSA coating and local changes in surface roughness and defects on the 3D printed resin surface. We would expect to see some variability of contact angle within the same part due to 3D printing resolution, but we expect intra-part variability to be smaller than variability between different parts. With both pinning geometries, the smaller angles pinned more consistently, while the larger angles (particularly those that are above where the hydrostatic pressure is greater than that of the threshold Laplace pinning pressure) failed to pin. In some cases, the average contact angle line predicted that pinning would occur, but it did not in all cases. However, when considering the range of possible contact angles, the lower bound lines on the graphs actually predicted failure, therefore we can assume the contact angle was smaller in those particular devices. A detailed description of these results is found in the Supplementary Text section 3. Additionally, we found that pipetting 1 µL of additional volume can change the bulging angle of the pinned fluid front’s vertical interface (see Supplementary Text section 4, Supplementary Table 2, and Supplementary Figs. 8-10 for detailed derivation). This phenomenon can help explain why experimentally we observed changes in successful pinning of the fluid front at different pipetted volumes. It should be noted that the theoretical conditions tested here are specific to parameters associated with a 5 mg/mL collagen solution. The specific cutoff values for pinning with the different features will change with different materials and with different channel and pinning geometries.

### Spatially heterogeneous engineered heart tissues (EHTs) generated with STOMP model cardiac fibrotic pathology

Fibrotic remodeling is an innate component of almost all acute and chronic cardiac pathologies, including hypertension, myocardial infarction, inherited cardiomyopathies, and end-stage heart failure[70]. This remodeling is characterized by an increase in the myocardial extracellular matrix, proliferation of stromal cells, and often a loss of cardiomyocytes[71]. Fibrotic progression is typically associated with an enhanced risk of arrhythmias and cardiac dysfunction[72]. Despite the near ubiquitous presence of fibrotic progression in cardiac disease, there are currently no clinical therapies available that directly address fibrosis[73]. While numerous investigators have developed various engineered heart tissue (EHT) platforms to model cardiac fibrosis, these systems are primarily homogeneous in cell type and ECM composition and fail to recreate the focal remodeling seen in vivo. Currently, the most advanced systems for modeling cardiac fibrosis using hiPSC-derived cardiomyocytes consist of a cardiac microtissue model utilizing a laser induced injury[74] and the Biowire II platform with two heteropolar healthy and fibrotic myocardial regions[75].

Using STOMP, we sought to develop an in vitro model of a cardiac tissue with a fibrotic interface, by creating spatially heterogeneous EHTs with a middle “fibrotic” region containing a localized excess of stromal cells and collagen, and deficit of cardiomyocytes, bordered on both sides by a “healthy” myocardial section (Figure 3a). Stromal cells used in the outer or middle regions were fluorescent HS5-GFP or HS5-mCherry cells, respectively, to delineate the regional boundaries and HS27A stromal cells were seeded throughout all regions (Figure 3a-b). To measure functional outputs of the patterned EHTs, both the control and fibrotic tissues were paced under 1.5 Hz frequency, resulting in the average twitch force waveforms seen in Figure 3c. Both control and fibrotic EHTs formed effective cardiac syncytia, as shown by their ability to follow the pacing frequency (Figure 3d). We next looked at the unpaced, spontaneous beat frequency of the EHTs, representative of a resting heart rate. Compared to control tissues, fibrotic tissues showed an elevated beat frequency, a response similar to that seen in patients with heart failure downstream of cardiomyopathy[76–78] (Figure 3e). Additionally, measurement of tissue contractions revealed that fibrotic EHTs presented with altered kinetics as they returned to diastole, or the phase where the heart chambers relax and refill with blood in preparation for the next contraction. This diastolic alteration in fibrotic EHTs was seen as a faster time to 50% relaxation (Figure 3f), but this difference is no longer evident at 90% relaxation (Figure 3g), suggesting that the fibrotic heart tissues initially relaxed faster but then slowed relaxation, compared to the control tissues. In patients with dilated cardiomyopathy, total relaxation time tends to be increased[79], therefore it is notable that in our model time to 50% relaxation was decreased. The increased stromal cells in the fibrotic tissue model may also partially explain the quicker time to 50% relaxation, a result that has been noted in previous in vitro models[80]. Finally, despite no differences in the twitch force (Figure 3h), specific force (Figure 3i), or shortening velocity (Figure 3j) compared to control tissues, fibrotic EHTs did display altered systole, or the contractile phase of cardiac cycle, as shown by a lower time to peak (Figure 3k).

**Figure 3.**
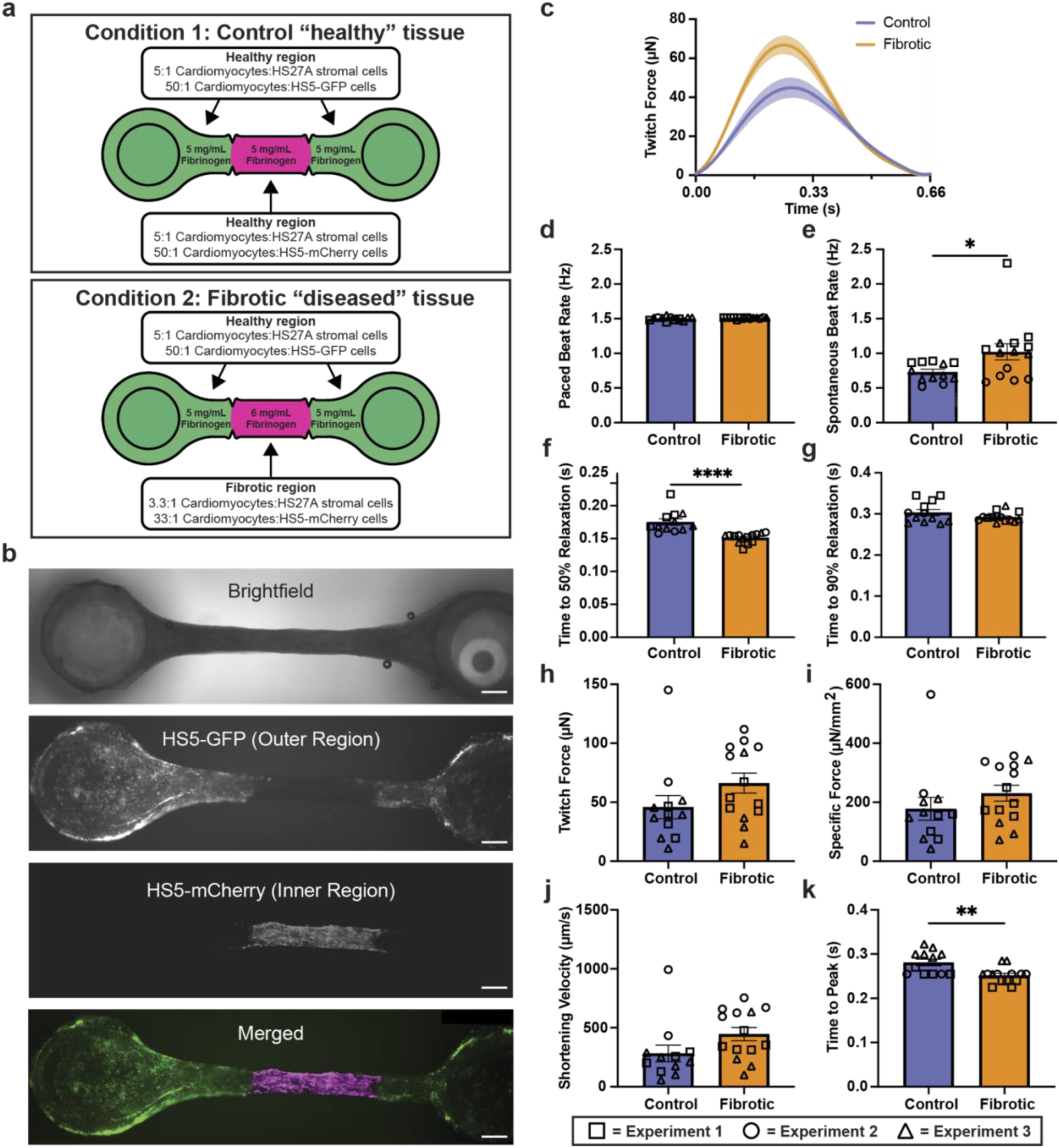
Patterned engineered heart tissues (EHTs). a) Conditions tested for modeling a fibrotic region in an EHT using STOMP, where the middle region has a higher fibrin content and lower cardiomyocyte density than the regions at either end. (a) Representative brightfield and fluorescent images of a control EHT seeded with HS5-GFP human bone marrow stromal cells (green) on the two outer regions near the flexible and rigid posts, and HS5-mCherry cells (magenta) in the center region. All scale bars are 500 μm. Full video of an unpaced EHT beating can be seen in Supplementary Movie 6 (GFP channel) and Movie 7 (mCherry channel). (c) Average twitch force traces of patterned control and fibrotic EHTs under 1.5 Hz pacing. (d) Paced beat rate of patterned control and fibrotic EHTs. (e) Spontaneous unpaced beat rate of patterned control and fibrotic EHTs. While all EHTs developed effective electromechanical coupling and were able to follow a pacing frequency of 1.5 Hz, fibrotic EHTs showed an elevated spontaneous beat rate when pacing was not applied. Diastolic function of control and fibrotic EHTs under 1.5 Hz pacing for (f) time to 50% relaxation and (g) time to 90% relaxation. Fibrotic EHTs show a delay in time to 50% relaxation and no change in the time to 90% relaxation. Systolic function of control and fibrotic EHTs under 1.5 Hz pacing, where fibrotic EHTs showed no differences in (h) maximum twitch force, (i) specific force, or (j) shortening velocity, but they did have a reduced (k) time to peak as compared to controls. Each shape (triangle, circle, square) represents an independent experiment for control EHTs (n = 12) and fibrotic EHTs (n = 14). Each data point is a separate tissue, with lines representing mean ± SEM. Statistical analysis was performed using unpaired t-test with two tails. *p≤0.05, **p≤ 0.01, ****≤0.0001.

Here we were able to demonstrate a defined border region between fibrotic and non-fibrotic tissue in an EHT model with STOMP. The ability to create this border region using STOMP will allow for further investigation of cardiac fibrotic pathology in longer-term cultures. In particular, having a healthy and fibrotic region within a single tissue will allow for investigation into pathological cardiac remodeling. However, further studies are required to validate STOMP as a platform for recapitulating pathological cardiac remodeling, such as investigations into cell-cell electrical and morphological couplings at the interface.

### STOMP enables a multi-tissue periodontal model with distinct regions and cellular enthesis

Periodontal ligament (PDL) attachment to alveolar bone is a key structural element that determines PDL function and behavior. The bone-ligament junction, or enthesis, in periodontal tissue constitutes a critical transition zone for stress distribution and functional adaptation to repeated cyclic mechanical loads that arise from oral functions like chewing. We have previously developed a 3D periodontal tissue construct (PTC) to investigate biomechanical behavior of PDL under tensile loads, however it is constituted of only PDL cells and lacks spatial patterning with different cell types[81]. Because the PTC model lacks the inclusion of the enthesis region, further investigations into periodontal health, disease, and regeneration at this interface are limited[82–84]. Here, we use STOMP to create a PTC model that incorporates spatially patterned regions of mineralization.

Specifically, we patterned osteogenic PDL cells in a collagen hydrogel on the outer regions connected by PDL cells in the center (referred to as OPO), mimicking the bone-PDL-bone arrangement found in vivo (Figure 4a). OPO PTCs were found to maintain these two cell populations in their specific locations as seen in bright field imaging (Figure 4b) along with corresponding spatial deposition of calcium deposits stained with alizarin red in the osteogenic outer regions (Figure 4c). Importantly, the PTCs had a distinct suture-like demarcation where the pinning features were located, transitioning from one tissue type to another. This border region indicates formation of a cellular enthesis representative of the fibrous union between PDL and alveolar bone in vivo[85].

**Figure 4.**
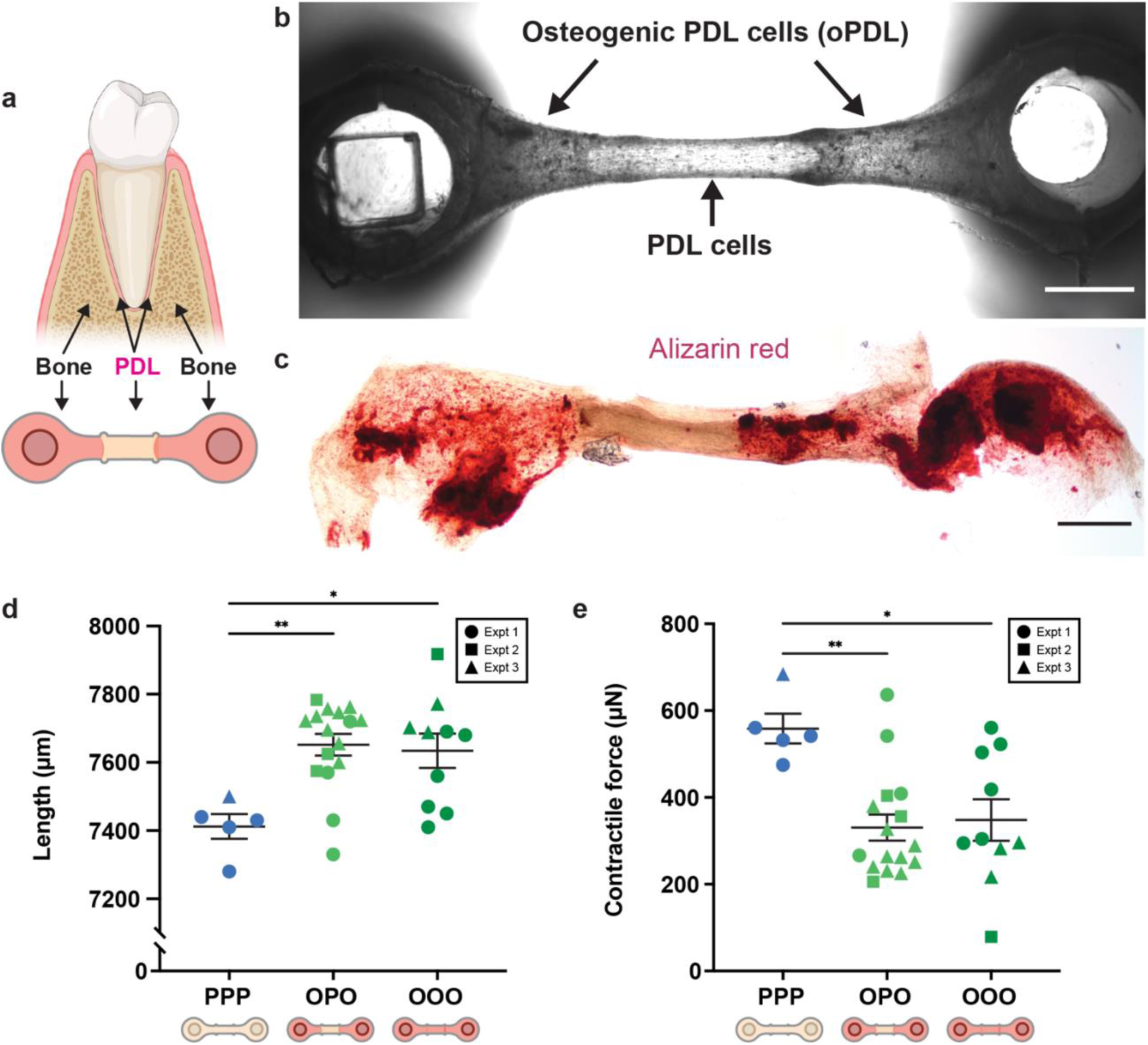
Patterned periodontal tissue constructs (PTCs) with a bone-periodontal ligament (PDL) border region. a) Visual representation of the bone-PDL junction observed in the human tooth with corresponding schematic of how this junction was patterned using STOMP. b) Representative brightfield image of the cellular entheses between osteogenic PDL cells (oPDL) in the outer regions and PDL cells in the inner region. Scale bar is 1 mm. c) Representative brightfield image of alizarin red assay showing mineralized calcium deposits (red) located in the outer regions containing oPDL cells. Scale bar is 1 mm. d) Final tissue length measurements of patterned oPDL-PDL-oPDL (OPO) tissues and control all periodontal (PPP) and all osteogenic PDL (OOO) tissues. e) Contractile force measurements of patterned OPO tissues and control PPP and OOO tissues. OPO tissues have contractile forces that are similar to OOO tissues and significantly lower than PPP tissues. Each shape (circle, square, triangle) represents an independent experiment for control PPP (n = 5), control OOO (n=10), and patterned OPO (n=16) tissues. Each data point is a separate tissue, with lines representing mean ± SEM. Statistical analysis was performed using a one-way ANOVA with Tukey’s multiple comparisons *post hoc* test. *p≤0.05, **p≤0.01.

Due to the suspended configuration, we were able to measure contractile forces in the patterned OPO, and the tissues containing only PDL cells (referred to as PPP) or only osteogenic PDL cells (referred to as OOO). Cells in the PTCs contract the ECM, pulling the flexible post towards the center. Patterned OPO tissues exhibited similar contractile forces to the tissues patterned solely with osteogenic PDL cells (OOO) (Figure 4e). However, both the OPO and OOO tissues exhibited significantly lower contractility and greater tissue length as compared to tissues patterned solely with periodontal ligament cells (PPP) (Figure 4d-e). This data indicates a change in biomechanical properties of tissues based on the function of constituent cells within the patterned regions. This observation is a valuable property recapitulated by PTCs made using STOMP, which can facilitate exploration of the biomechanical behaviors within and interactions between the multiple cell-layers of the PDL.

### STOMP enables versatile tissue geometries and expanded patterning capabilities from suspended cores to patterning non-compactable cell-ECM combinations and designer materials

In all patterning implementations described thus far, we show a geometry where we are changing the region along one axis. However, we expand upon the geometric versatility capabilities of STOMP by creating a tissue with an inner core region fully surrounded by a second outer region, which enables patterning along the z-axis in addition to the x-y axes (Figure 5a-c). The process for generating a core region is shown in Figure 5a. After patterning an inner core region with any configuration of STOMP (e.g., one, two, or three regions), a second patterning device is added that has an open channel that is larger in length, width, and height than that of the first patterning device; this outer region can then be patterned around the first hydrogel, generating a suspended tissue that is completely encapsulated by another cell-laden hydrogel (Figure 5a). We demonstrate this ability with an inner core composed of a single region of 3T3 cells dyed by CellTracker Red CMTPX laden in fibrin and an outer core composed of 3T3 cells dyed by CellTracker Green CMFDA laden in fibrin (Figure 5b). Further, we show multiple combinations of the inner core containing one, two, and three-patterning regions that are then completely encapsulated by an outer region with colored agarose (Figure 5c). In future implementations, the outer region could also contain multiple regions by adding pinning features to the outer region patterning device.

In addition to expanding upon the geometric capabilities of STOMP, we also implement strategies to expand the types of hydrogels that can be used in the STOMP system. The requirement of the tissue pulling away from the channel walls of the patterning device limits STOMP to cell-ECM combinations that allow for sufficient tissue compaction or materials that do not adhere strongly to channel walls (such as in the case for agarose). To address this limitation, we designed a STOMP device with a v-shaped channel carved into the walls of the patterning channel (Figure 5d). We pipetted a degradable hydrogel into this channel (for demonstration we use a sortase-degradable PEG, although any degradable hydrogel can be used); after gelation, this degradable hydrogel becomes the STOMP channel walls. We could then pipette a second solution into the middle tissue region. After patterning, the channel wall can be exogenously triggered to degrade, thus allowing for facile release of tissue from the patterning device without a requirement for tissue compaction. A schematic for the patterning process of the degradable “wall” region and tissue region is shown in Supplementary Figure 13, and a still image from a video with colored agarose for visualization is shown in Figure 5e. Using this method, we successfully patterned a second PEG-based designer material laden with 3T3 fibroblast cells that otherwise cannot be removed from the STOMP device after patterning (Figure 5f). After the cell-ECM PEG tissue region was gelled, the entire assembly was placed in a solution containing sortase specifically engineered to recognize and cleave the cross-linker in the PEG pipetted into the v-shaped channel walls. After degradation of the PEG hydrogel in the v-shaped channel, the tissue patterned in the center tissue region was released as the channel “walls” degraded. Adding the new degradable wall feature allows STOMP to be used for synthetic materials that do not compact readily (e.g., stimuli responsive or “4D” PEG-based materials that can be patterned in space and triggered in time)[57,86–88] or for cell types that do not readily remodel ECM and compact (e.g., nerve cells in a neuromuscular junction). This addition extends STOMP to virtually any tissue or material, opening a frontier in 4D suspended multiregion tissues.

**Figure 5.**
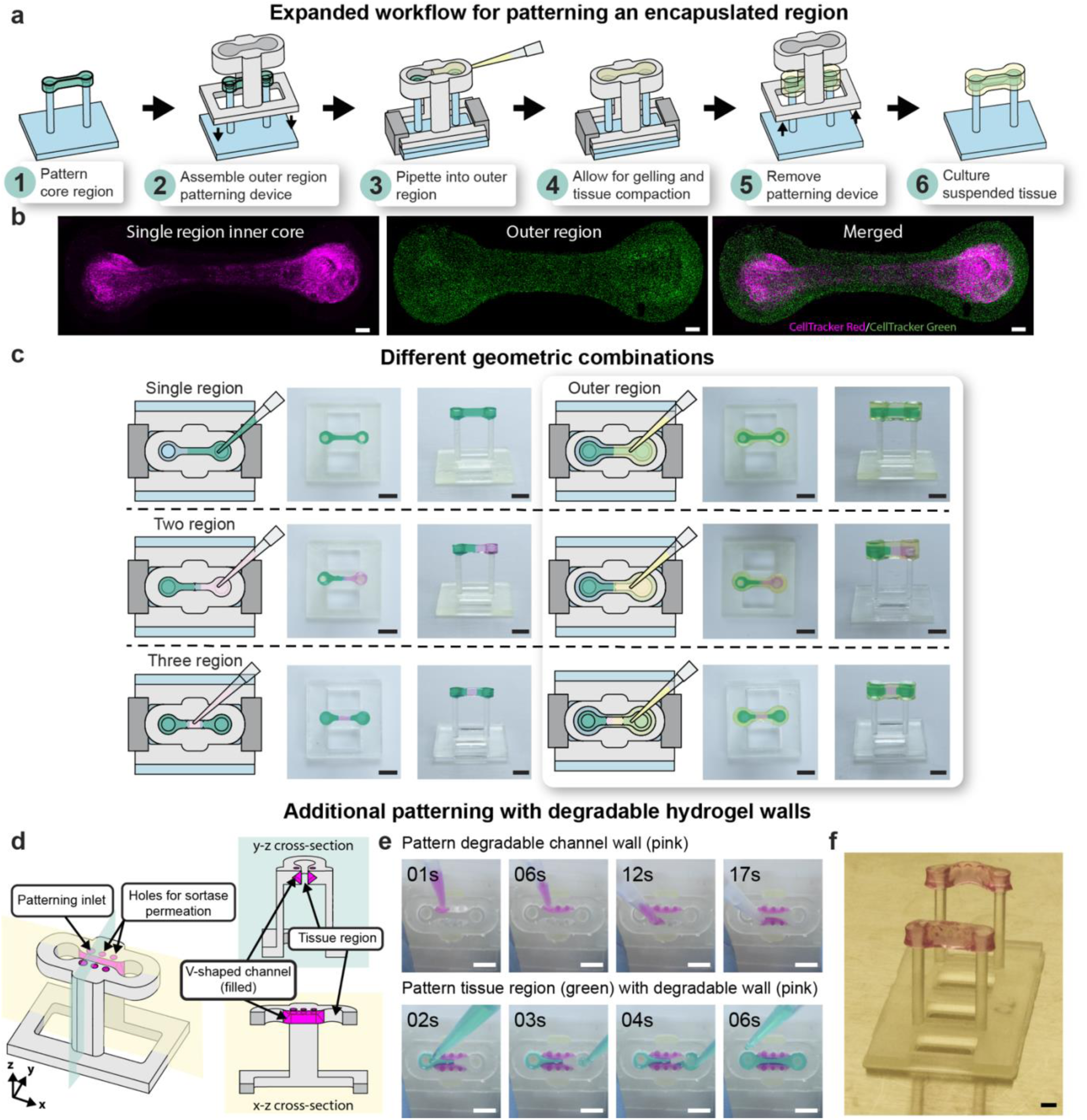
Expansive geometric and patterning capabilities of STOMP. a) Workflow of patterning an inner core region completely encapsulated by an outer region, thus patterning suspended tissues in the z-direction. b) Representative fluorescent images of patterned 3T3 mouse fibroblast cells laden in fibrin to generate a suspended tissue with an inner region and outer region. The inner region tissue was generated with the single region STOMP configuration; 3T3 cells were dyed by CellTracker Red CMTPX (magenta). The outer region contains 3T3 cells dyed by CellTracker Green CMFDA (green). All scale bars are 500 μm. c) Demonstration of patterning single region, two region, and three region suspended tissue constructs using colored 1.5% agarose; additionally, these patterned tissues can be completely encapsulated by placing a patterning device larger in width, height, and length around the previously patterned tissue to generate an encapsulated core. All scale bars are 4 mm. d) Schematic of a STOMP patterning device containing a V-shaped channel within the walls of the patterning device tissue region’s open channel. This geometry can be used to pattern non-compactable synthetic hydrogels, such as poly-ethylene glycol (PEG), where a degradable PEG hydrogel can be patterned in the V-shaped channel and degraded after patterning in the tissue region. e) Representative video still images of colored agarose patterned in the v-shaped channel wall region (pink hydrogel) and tissue region (green hydrogel). All scale bars are 2 mm. Full patterning Video can be seen in Supplementary Video 5. f) Representative image of resulting suspended PEG hydrogel laden with 3T3 mouse fibroblast cells. Scale bar is 2 mm.

## DISCUSSION

The STOMP platform is an open microfluidic approach for generating spatially controlled suspended tissue models. Precise spatial patterning can be achieved in the STOMP system via capillary pinning in the x-y direction (i.e., within a single tissue) or along the z-direction (i.e., generating encapsulated tissue structures). With STOMP we are able to achieve a wide range of geometrical combinations, including structures that are difficult to pattern with existing fabrication methods such as the fully encapsulated suspended tissue. In this initial proof-of-concept demonstration of STOMP, we show that STOMP can be used to model tissue interfaces (e.g., periodontal ligament-bone) and intra-tissue heterogeneity (e.g., fibrotic and healthy tissue co-culture), thus establishing the versatility and board applicability of STOMP for modeling complex multiregional suspended architectures. Lastly, we demonstrated the use of degradable PEG-based hydrogels, a material that may not meet requirements for other 3D suspended patterning methods. The ability to pattern stimuli-responsive synthetic hydrogels opens up a fourth dimension (i.e., time) of control to the suspended patterned tissues, allowing for STOMP to be used in 4D suspended tissue engineering applications. STOMP is designed to complement existing post-based suspended culture systems; its strength lies in its ability to integrate easily into these tissue fabrication workflows while adding regional control via open microfluidic principles.

Existing post-based suspended tissues protocols utilize a casting method, where a cell-ECM liquid precursor solution is pipetted into a mold made of agarose or PDMS that surrounds the suspended posts[17,18,26,27]. Because there are no features to compartmentalize these molds, regional control has not previously been demonstrated, limiting studies to a homogenous singular region. Other methods, such as 3D bioprinting and photopatterning, allow for the additional fabrication complexity of patterning multiple regions within a single tissue. However, these methods typically require highly viscous, extrudable materials or photoactive moieties, making it difficult to pattern unmodified native ECMs. Recently, there have been advancements in 3D bioprinting techniques to mitigate these limitations on materials by printing cell-laden hydrogels into a sacrificial gelatin suspension bath[29,32,33]. These approaches enable the 3D printing and spatial patterning of natural polymers, such as alginate, collagen, fibrin, and hyaluronic acid. While suspended biofabrication methods allow for the generation of large-scale structures (millimeter- to centimeter-scale), these methods typically require large volumes (milliliter to tens of milliliters) of hydrogel material—both for the sacrificial support and the final suspended structure—which can limit the use of precious biomaterials. Additionally, these techniques often rely on custom-engineered 3D bioprinting systems, restricting scalability and accessibility. In comparison to other suspended biofabrication methods, STOMP offers finer volumetric control at smaller scales (30 µL - 100 µL), making it a more efficient and scalable alternative for applications requiring precise hydrogel patterning without excessive material consumption. Taken together, STOMP fills an important niche for simple, microliter-scale, and multi-region in vitro modeling; other methods such as 3D bioprinting will remain important for large-scale, high resolution organ modeling.

In the STOMP system, material limitation is governed by the removal process. While surface tension-driven flow allows virtually any hydrogel that transitions from a liquid to a solid state to flow in the STOMP channel, some hydrogel materials can adhere to the STOMP channel during the removal process. Therefore, when adapting the STOMP workflow to a new tissue model, it is important to consider material and tissue type compatibility with the STOMP device. For example, we experienced significant challenges while using the PEG-based degradable hydrogels in the original implementation of STOMP (i.e., without the degradable walls), because the material did not allow for sufficient compaction of the tissue to facilitate successful removal of the STOMP device. This issue would be similar in other cases using compactable ECMs (e.g., fibrin or collagen) with a cell line that typically does not remodel ECM or contract (e.g., neuronal cells without supporting fibroblasts). While we do present a work-around to this issue with degradable channel walls, the degradable walls add complexity, require the use of more expensive degradable materials, and limit the addition of pinning features due to the difficulty of patterning.

Once a hydrogel is determined to be compatible with the removal of the STOMP device, there are two additional practical considerations for patterning multi-region tissues: first, determining the best pinning feature shape and angle and second, determining if two regions at the defined interface can integrate to form a contiguous tissue. In this work, we provide general equations for users to insert their specific hydrogel density and contact angle to determine the relationship between hydrostatic pressure and maximum Laplace pinning pressure in the STOMP device (Eq. 5, 7, and 8). To demonstrate the workflow, we specifically calculate these relationships using the density and contact angle of a 5 mg/mL precursor collagen solution on a 1% BSA treated 3D printed surface (the same treatment done to the STOMP device before patterning), allowing us to determine the best pinning feature and angle for workflows involving collagen (i.e., 90° concave “cavity” pinning features). However, when working with fibrin, we found that the convex “vampire” pinning features resulted in better patterning (i.e., pinning and subsequent de-pinning to connect two regions at the defined interface). If using a different hydrogel other than collagen and fibrin, the user would benefit from utilizing the equations detailed in the work as well as experimentally testing the best pinning feature shape and angle that will result in the most reproducible patterning for their tissue. After optimizing the pinning geometry, it must next be determined if two regions at an interface can form a functional, contiguous tissue. Two regions will more easily form a contiguous tissue if each region utilizes similar ECM and cell compositions, such as two separate cell types with similar propensity to contract ECM or two different ECMs (e.g., collagen and Matrigel) that have similar components and can bind together. However, collagen and fibrin are two common ECMs for 3D tissue modeling, and while these two ECMs are compositionally different, in preliminary experiments we have patterned cell-laden tissues with a collagen-fibrin interface, which successfully formed a contiguous tissue. This example illustrates the importance of testing each new ECM and cell-type for patterning compatibility. If the desired interface requires materials that are incompatible, STOMP would not be a suitable platform.

In addition to material considerations, there are also dimensional considerations when adapting the STOMP platform to specific experimental needs. It may be desired to have a smaller tissue volume (micrometer-wide channels with millimeter-scale separation between the two posts) when using precious cells or materials or increasing the throughput of an experiment. In this case, the principles of surface tension-driven flow would make flow in the STOMP system (i.e., a channel without a ceiling or floor) more desirable since there would be less of an effect due to gravity at smaller scales, resulting in less bulging of the liquid underneath the channel. Since the hydrostatic pressure at the bottom of the channel would be decreased in a smaller channel design (e. g., a channel 0.2 mm in width and 1.0 mm in height) compared to a larger channel design (e.g., a channel 0.8 mm in width and 3.5 mm in height as detailed in this work), the pinning in smaller channel dimensions would be more stable. Therefore, the smaller channels would also be less sensitive to changes in volume due to increased maximum Laplace pinning pressure (Eq. 5 and Eq. 7) and decreased hydrostatic pressure (Eq. 8) observed in channels with decreased w and h values. However, at smaller dimensions, the tolerance of the 3D printer used will have a greater effect on the channel dimensions and therefore also on the total volume that would completely fill the channel; a tolerance of 20 µm would have a larger effect on the volume of a channel that is 200 µm wide than a channel that is 800 µm wide. This potential variability could be mitigated by using a higher resolution printer with tighter tolerances or with a different fabrication technique (e.g., CNC milling or injection molding). Smaller channel length dimensions also limit the number of different regions that can be generated within a single tissue since it is practically challenging to pipette accurate volumes into small regions.

For larger tissue volumes (millimeter-wide channels with centimeter-scale separation between the two posts), which could be desired for longer-term tissue cultures that are highly contractile and more prone to breaking over time, there are additional limitations and challenges. When increasing the width of the channel, the height of the channel must also be increased to fulfill the equation for spontaneous capillary flow in the STOMP system (Eq. 2). Increasing the volume will cause a greater effect due to gravity which will exacerbate non-uniform filling of the channel cross-section, with more liquid bulging at the bottom of the channel. Since hydrostatic pressure increases and maximum Laplace pinning pressure decreases in larger channels, pinning is less stable overall. Therefore, de-pinning events, which typically occur at the bottom of the channel where hydrostatic pressure is the greatest, in larger channels would be more sensitive to changes in volume pipetted.

While optimizing STOMP for a specific workflow requires careful consideration of materials, dimensions, and interface integration, once established, it enables seamless spatial patterning in a suspended system with just a few pipetting steps. This adaptability makes STOMP a powerful and flexible tool for advancing complex suspended tissue engineering applications. Our ability to pattern multiple regions within the tissue enhances integration of different cell types and ECMs, enabling complex studies of tissue interactions, maturation, and disease progression. These benefits can be combined with the suspended nature of the tissue, which provides axial stretching and mechanical cues, and the use of flexible PDMS posts for controlled manipulations and functional measurements. STOMP’s versatility in integrating different cell types and ECMs, coupled with the mechanical benefits of suspended tissues, opens new experimental possibilities and advances our understanding of tissue behavior in both healthy and diseased states.

## METHODS

### Device Design, Fabrication, and Sterilization

Post arrays made from polydimethylsiloxane (PDMS) were fabricated as described in Bielawski et al[59]. Briefly, uncured PDMS was poured into a four-part acrylic mold to fabricate an array that contains six pairs of posts and that fits in a row of a 24-well plate. Glass capillary tubes were cut to the appropriate length and inserted into one side of the mold prior to curing such that each pair contained one rigid post. The posts were baked overnight at 65°C and then removed from the molds. Fabricated posts were 8 mm apart in spacing, 12 mm in length, and 1.5 mm in diameter with a cap structure (0.5 mm thick and 2.0 mm diameter) to aid in attachment of the cell-laden hydrogel of interest.

For STOMP, the open microfluidic patterning device and clips were designed in Solidworks and 3D printed out of Formlabs clear resin V4 using a Form 3B+ 3D printer (Formlabs Inc.). Dimensions of the open channel were designed to fit around the PDMS posts; dimensions of the three open channel designs used in this work are depicted in Supplementary Figure 1. Clips were designed to hold the patterning device in place during the tissue patterning process. The patterning devices and clips were cleaned in two separate FormWash units (Formlabs Inc.) with isopropyl alcohol (IPA) for 20 minutes and then another 10 minutes to remove excess uncured resin. The devices were dried with compressed air and cured under UV light at 60°C for 15 minutes in a FormCure (Formlabs Inc.).

To sterilize the PDMS posts, patterning devices, and clips for cell culture, all parts were first placed within a biosafety cabinet and UV sterilized for 20 minutes. The PDMS posts were additionally sterilized with 70% ethanol for five minutes by placing the tips of the posts in a 24-well plate, where each well contained 2 mL of 70% ethanol. The PDMS posts were then rinsed with sterile DI water twice for five minutes each.

After sterilization, the patterning devices that were to be used with a cell-laden hydrogel or ECM were incubated in a solution of 1% bovine serum albumin (BSA) for 1 hour at room temperature. After incubation, the 1% BSA solution was aspirated, and the patterning devices were allowed to fully dry prior to assembling the suspended tissue open microfluidic patterning devices. The open microfluidic patterning device was placed such that it surrounded the ends of each post, and the clips were used to hold the patterning device in place at the base of the PDMS post array.

### Experimental Pinning Analysis

#### Measuring contact angle of 5 mg/mL collagen on BSA treated resin

3D printed resin squares were utilized to provide a flat surface for contact angle measurements. Each square was treated with 1% BSA for 1 hour at room temperature. A Drop Shape Analyzer DSA25 (Krüss) was utilized on the sessile drop mode to measure contact angle. A SY3601 disposable 1 mL syringe with Luer connector (Krüss) was filled with DI water and then fitted with a NE94 steel needle with a 0.51 mm diameter (Krüss). The DSA25 was calibrated to the image of the needle. A collagen solution was prepared by mixing a stock of rat tail collagen type I (Corning) in 0.02 N acetic acid with sterile deionized water, 10X PBS, and 1N NaOH to achieve a final collagen density of 5 mg/mL and kept on ice to prevent gelation. A BSA treated resin square was placed on the platform below the needle and a 2 µL droplet of the collagen solution was dispensed via pipetting. The contact angle of each drop was measured using multiple images of the collagen sessile drop at the points of intersection (three-phase contact points) between the drop contour and the projection of the surface baseline using ADVANCE software (Krüss). Twenty different drops were measured on 20 different BSA treated resin squares (one drop per resin square); the mean contact angle and standard deviation of each drop were calculated by the Krüss Drop Shape Analyzer based on the number of measured technical replicates, which are multiple images of the same droplet (see raw data in Supplementary Table 1). The experiment number indicated in Table SI (Experiment 1-3) indicates which experiment number a particular resin part belongs to (i.e., we performed these contact angle measurements on three different days with different batches of 3D printed surfaces and 1% BSA solutions). The overall average contact angle of the 20 drops (see raw data in Supplementary Table 1). The average contact angle and the minimum and maximum contact angle measurement were used for theoretical modeling of the pinning of collagen in STOMP.

#### Measuring success or failure of collagen pinning in STOMP system

The STOMP devices with a channel height of 3.5 mm and a width of 1.2 mm were used for the pinning experiment. All STOMP patterning devices were treated with 1% BSA for 1 hour. A collagen precursor hydrogel solution was prepared as described above. Pipettes were set to 23, 25, 26 or 27 µL to dispense collagen precursor slowly at the circular outer section of the STOMP channel, stopping at the first stop of the pipette to avoid bubble introduction. The pipette set to 23, 25, 26, and 27 µL dispensed actual volumes of 20, 22, 23, and 24 µL, respectively, as measured by weight in triplicate and using the density of 5 mg/mL precursor collagen solution (958 kg/m3). Throughout the text, the actual volumes dispensed for the pinning experiments will be referenced. The flow was observed for 10s. If the flow moved past the pinning feature, then the replicate was considered a “failure” of pinning. If the fluid front remained pinned at the pinning feature at the end of 10s then it was deemed a “success” of pinning. Each pinning feature, angle, and volume was done in triplicate. Figure 2e-f shows representative images of examples of successful pinning and no pinning in both pinning feature designs; the volume of collagen used in these pictures was 23 μL. The complete dataset is visualized in Supplementary Figure 2 and representative videos shown in Figure 2e-f are found in Supplementary Movies 2-5.

### Cell Culture and Maintenance

#### 3T3 mouse fibroblasts

Initial optimization experiments were conducted with NIH/3T3 mouse embryonic fibroblast cell line obtained from ATCC. The cells were maintained in a tissue culture flask containing Dulbecco’s Modified Eagle Medium (DMEM) supplemented with 10% fetal bovine serum (FBS) and 1% penicillin-streptomycin at 37°C, 5% CO2. Culture medium was changed every other day until cells reached around 80-90% confluency, whereupon the culture was rinsed with 1X PBS, followed by addition of TrypLE Express (Gibco). After incubation for 3 minutes at 37°C, the TrypLE was inactivated by diluting with cell culture medium. The fluid volume was centrifuged at 300 RCF for 3 minutes. The cells were resuspended in cell culture medium for further passaging or used for hydrogel encapsulation, as described below.

#### Human induced pluripotent stem cell (hiPSC) derived cardiomyocytes

WTC11 hiPSCs were differentiated using a modified small molecule Wnt-modulating protocol as previously described[26,89–91]. Cells were seeded at a density of 1.6-5.8 x 104 cells/cm2 on plates coated with a 1:30 dilution of Matrigel in mTeSR+ (STEMCELL Technologies) supplemented with 100 U/mL penicillin-streptomycin and 5 µM ROCK inhibitor Y-27632. Upon reaching 50-80% confluency (Day-1), cells were washed with 1X PBS and the media was changed to mTeSR+ supplemented with 1 µM CHIR-99021 (Cayman Chemical). After 24 hours (Day 0), cells were washed with 1X PBS and media was replaced with RBA (RPMI 1640, 500 µg/mL BSA, 220 µg/mL ascorbic acid, 100 U/mL penicillin-streptomycin) supplemented with 3-5 µM CHIR-99021. After 48 hours (Day 2), cells were washed with 1X PBS and media was replaced with RBA supplemented with 2 µM Wnt-C59 (SelleckChem). After 48 hours (Day 4), cells were washed with 1X PBS and the media was replaced with RBA without supplementation. After 48 hours (Day 6), cells were washed with 1X PBS and media was replaced with cardiomyocyte media (RPMI 1640, B27 plus insulin, 100 U/mL penicillin-streptomycin) and replaced every other day until cells began beating on Day 10-11. hiPSC-CMs were then washed with 1X PBS and metabolically enriched by culturing in DMEM without glucose supplemented with 4 mM sodium L-lactate for 4 days. hiPSC-CMs were then returned to cardiomyocyte media for 48 hours. hiPSC-CMs were then replated at a density of 1.5-1.8 x 105 cells/cm2 on plates coated in a 1:60 dilution of Matrigel in cardiomyocyte media supplemented with 5% FBS and 5 µM ROCK inhibitor Y-27632. After 24 hours, the media was replaced with standard cardiomyocyte media. After 24 hours, hiPSC-CMs washed with 1X PBS and again metabolically enriched by culturing in DMEM without glucose supplemented with 4 mM sodium L-lactate for 4 days. hiPSC-CMs were then returned to cardiomyocyte media with the media replaced every other day until Day 35-40 when used for EHT casting.

WTC11 hiPSCs were cultured in mTeSR+ supplemented with 100 U/mL penicillin-streptomycin on plates coated with a 1:30 dilution of Matrigel. Cells were fed every other day. Media was supplemented with 5 µM ROCK inhibitor Y-27632 for the first 24 hours after passing.

#### Human periodontal ligament (PDL) cells

PDL cells were harvested from the roots of healthy premolars extracted for orthodontic purposes, using pre-established protocols[92]. Briefly, PDL tissue was gently dissected from the middle third of the tooth root surface in a culture dish containing wash buffer [Hanks’ balanced salt solution (HBSS; Gibco) supplemented with 5% (v/v) FBS (HyClone) and 50 U/mL penicillin and 50 μg/mL streptomycin]. The suspension was centrifuged at 400 RCF for 10 min at 4 oC. After aspirating the supernatant, PDL tissue was digested in collagenase I (3 mg/mL, Sigma-Aldrich) and dispase II (4 mg/mL; Sigma-Aldrich) for 1 hour at 37 oC. After digestion, the suspension was passed through a 70 μm strainer to obtain a suspension of isolated cells.

The population of cells was expanded by seeding them in 24-well plates. Cells were grown in DMEM supplemented with 10% FBS (HyClone) and 1% penicillin-streptomycin at 37 oC with 5% CO2. On reaching confluence, cells were washed with PBS, trypsinized with 0.25% trypsin-EDTA (Gibco), and passaged progressively into T-75 and T-175 flasks. Cells at passage numbers 1 to 3 were stored in freezing media Cryostor (1 mL per 10 million cells) in a liquid nitrogen freezer then thawed, plated, and expanded as needed in T-75 and T-175 flasks. PDL cells between passage numbers 3 and 8 were used for the patterning experiments.

#### Osteogenic differentiation of PDL cells

To induce osteogenic differentiation, a sub-population of PDL cells between passages 3 to 6 was transferred to a T-75 flask and cultured in DMEM media with 10% FBS and 1% penicillin-streptomycin. The cells were left to grow up to 80-90% confluence with culture medium changes every 3 days before adding osteogenic induction medium (culture medium supplemented with 50 μM ascorbic acid-2-phosphate, 10 mM β-glycerophosphate, and 10 nM dexamethasone) for 2 weeks. After 2 weeks, these cells were trypsinized and lifted to be used for hydrogel encapsulation.

### STOMP Tissue Preparation

#### Patterning with agarose

Low gelling temperature agarose (MilliporeSigma) was dissolved in deionized water to a concentration of 15 mg/mL and dyed for visual aid with food dye (Spice Supreme) or India ink (Dr. Ph. Martin’s). Prior to patterning, the agarose solution was heated to a liquid state (95°C) and then pipetted into STOMP devices. To generate Figure 1g-h, agarose was pipetted into a STOMP device with a channel height of 2 mm and width of 1 mm and the three region 60° convex “vampire” pinning features. For generating agarose structures seen in Figure 5, the STOMP devices used to generate the inner core had a channel height of 2 mm and width of 1 mm and either the two or three region 20° convex “vampire” pinning features. For the outer core patterning, a STOMP device with a channel height of 4 mm and width of 1.9 mm was used. Agarose structures were allowed to gel at room temperature for 5- 10 minutes. All agarose structures used 1.5% wt/v gel.

#### Patterning 3T3 mouse fibroblast cells with fibrin

Fibrinogen stock solutions were prepared at 50 mg/mL by dissolving 250 mg of powdered fibrinogen from human plasma (Sigma-Aldrich) in 5 mL of warmed 0.9% NaCl in deionized water. The fibrinogen solution was then filter-sterilized with a 0.22 mm syringe filter. Thrombin stock solutions were prepared at 100 U/mL by dissolving 100 units of powder thrombin from human plasma (Sigma-Aldrich) in 1 mL of warmed 0.9% NaCl in deionized water. Fibrinogen and thrombin stock solutions were aliquoted and stored at −20°C until ready for use.

After dissociating the 3T3 cells, two aliquots of the cells were dyed by either CellTracker Green CMFDA or CellTracker Red CMTPX at a final concentration of 25 μM in DMEM for 30 minutes at 37°C. After dying, the cells were pelleted and resuspended at a final concentration of 5 x 106 cells/mL in DMEM with 5 mg/mL bovine fibrinogen and 3 U/mL thrombin. For patterning in Figure 1f, the STOMP device with a channel height of 3.5 mm and width of 1.2 mm and the three region 60° convex “vampire” pinning features was used. For patterning in Fig 5b, the inner core was patterned with a STOMP device containing a channel height of 2 mm and width of 1 mm and the outer core was patterned with a STOMP device containing a channel height of 4 mm and width of 1.9 mm. After patterning, the 3D printed posts with the patterning device and clips assembled were placed in a 24-well plate and allowed to gel for 30 minutes at 37°C. After gelling, 2 mL of growth media supplemented with 5 mg/mL 6-aminocaproic acid (Sigma-Aldrich) was added to each well. After two days in culture, the STOMP devices were removed from the 3D printed posts and fresh media was added to the wells. Tissues were cultured for one day after the STOMP patterning device was removed then fixed with 4% paraformaldehyde (PFA) for one hour at 4°C. Tissues were rinsed twice with 1X PBS and stored at 4°C until ready for imaging. Tissue samples were imaged by placing the tissue directly on a coverslip without removing from the posts. All tissues were imaged on a Leica SP8 confocal microscope at 10X magnification.

#### Patterning engineered heart tissues (EHTs) with fibrin

Fibrinogen stock solutions were prepared at 50 mg/mL by dissolving 1 g of fibrinogen from bovine plasma (Sigma-Aldrich) in 20 mL of warmed cardiomyocyte media. Thrombin stock solutions were prepared at 100 U/mL by dissolving 1000 U of thrombin from bovine plasma (Sigma-Aldrich) in 10 mL of a 3:2 ratio of PBS to H20. EHTs were cast on PDMS posts using a modification of our previously described protocol[59,60,90]. For generating the EHTs, STOMP devices with a channel height of 3.5 mm and width of 1.2 mm and the three region 60° convex “vampire” pinning features were used. The two outer regions of all tissues were patterned using a solution of 27 µL volume consisting of 1.35 x 105 hiPSC-CMs (5 x 106 cells/mL), 2.7 x 104 HS27a human bone marrow stromal cells (1 x 106 cells/mL), and 2.7 x 103 HS5-GFP human bone marrow stromal cells (1 x 105 cells/mL) in cardiomyocyte media with 5 mg/mL bovine fibrinogen and 3 U/mL thrombin. The center region of control tissues was patterned using a solution of 12 µL volume consisting of 6.0 x 104 hiPSC-CMs (5 x 106 cells/mL), 1.2 x 104 HS27a human bone marrow stromal cells (1 x 106 cells/mL), and 1.2 x 103 HS5- mCherry human bone marrow stromal cells (1 x 105 cells/mL) in cardiomyocyte media with 5 mg/mL bovine fibrinogen and 3 U/mL thrombin. The center region of fibrotic tissues was patterned using a solution of 12 µL volume consisting of 4.8 x 104 hiPSC-CMs (4 x 106 cells/mL), 1.44 x 104 HS27a human bone marrow stromal cells (1.2 x 106 cells/mL), and 1.44 x 103 HS5-mCherry human bone marrow stromal cells (1.2 x 105 cells/mL) in cardiomyocyte media with 6 mg/mL bovine fibrinogen and 3.6 U/mL thrombin. After the tissues were patterned, the PDMS post array with the patterning device and clips assembled were placed in a 24-well plate and allowed to gel for 60 minutes at 37°C. After gelation, the tissues were transferred to wells containing cardiomyocyte media supplemented with 5% DMEM, 5% FBS, 1% non-essential amino acid (NEAA), and 5 mg/mL aminocaproic acid. After 48 hours, the media was replaced with cardiomyocyte media supplemented with 5% FBS, 1% NEAA supplement, and 5 mg/mL aminocaproic acid. After 24 hours, the patterning devices were removed from the EHTs and the tissues were transferred to EHT media (cardiomyocyte media supplemented with 5 mg/mL aminocaproic acid). EHT media was replaced every 2-3 days for 18 days until subsequent analysis.

#### Patterning periodontal tissue constructs (PTCs) with type I collagen

Following sterilization, PDMS posts were treated with 0.1% polyethylenimine (PEI) for 10 minutes and rinsed with sterile deionized water for 5 minutes. Next, the PDMS posts were treated for 30 minutes with 0.01% glutaraldehyde and then rinsed twice with sterile deionized water for 5 minutes each. For each treatment step, 2.5 mL/well of the respective solution was placed in a 24-well plate so that the tips of each PDMS post were in contact with the solution.

For generating PTCs, STOMP devices with a channel height of 2 mm and width of 1 mm and the three region 90° concave “cavity” pinning features were used. To pattern the tissues, two different cell-collagen mixtures were prepared with the PDL cells and osteogenic PDL (oPDL) cells. To prepare the cell-collagen mixtures, each cell type was mixed with type I rat tail collagen such that a 100 μL cell-collagen mixture was composed of 1:4 ratio of cell solution (20 μL culture media containing 300,000 cells) and collagen mixture [80 μL of 9 parts of collagen (4 mg/mL; Advanced Biomatrix) mixed with 1 part of neutralization solution (Advanced Biomatrix)]. Each cell-collagen mixed had a final concentration of 3×106 cells/mL.

First, 13 μL of the oPDL cell-collagen mixture was pipetted on one of the circular ends of the STOMP channel that surround the PDMS posts and then the other. Then, 3-4 μL of PDL cell-collagen mixture was pipetted in the center region of the STOMP channel. The PDMS posts, with the patterning rail and cell-ECM precursor solution, were then placed upside down in a 24-well plate. The assembly was transferred to the incubator at 37oC with 5% CO2. After 90 min, 2.5 mL of culture media was gently added to the wells containing the PTCs and then incubated again at 37 oC. After 48 hours, the clips and patterning rails were removed to reveal a suspended tissue between the PDMS posts. These patterned PTCs (referred to as OPO) were then transferred to a fresh 24-well plate with 2.5 mL/well of culture medium, which was changed every 2 to 3 days. Similarly, patterned control tissues consisting of oPDL cells in all 3 regions (referred to as OOO) and periodontal cells in all 3 regions (referred to as PPP) were also generated.

#### Patterning poly(ethylene glycol) (PEG) in modified STOMP

To pattern tissues with enzymatically degradable PEG as the ECM, we used PEG-tetraBCN (a four-arm PEG end-capped with reactive bicyclononyne groups) with two different sortase-degradable and bis(azide)-modified peptide crosslinkers. These materials are described in detail in previous work[57,86,93]. The synthesis of the PEG-tetraBCN and crosslinkers are described in detail in the Supplementary Methods sections 2-4. The LPESG crosslinker is cleaved by the eSrtA(4S9) sortase, and the LAETG is cleaved by the eSrtA(2A9) sortase. Briefly, PEG was prepared by combining the peptide crosslinker with a final concentration of 8mM, PEG-tetraBCN with a final concentration of 4mM. Components were vortexed together to ensure homogeneity. The PEG that served as the degradable “channel wall” was pipetted into the v-shaped channel. In this case, we used the eSrtA(2A9)-sensitive LAETG crosslinker in the v-shaped channel. There were no cells in the outside channel, therefore the PEG-BCN was suspended in 1X PBS. We used the eSrtA(4S9)-sensitive LPESG crosslinker in the tissue region that made up the final tissue, although we did not degrade this region. Because this region was laden with NIH 3T3 mouse fibroblasts at a final concentration of 5 x 106 cells/mL, RGDS peptide, the adhesion motif from fibronectin, was added at a final concentration of 1 mM, and the mixture was resuspended in cell culture media. For the tissue region, the height of the channel was 2 mm and the width of the channel (measured as the distance between the degradable channel walls) was 1 mm.

We observed that the PEG hydrogel made with the LAETG crosslinker would swell more when left in media than the PEG hydrogel made with the LPESG crosslinker. We therefore decided to use the LPESG crosslinker as the region that made up the final tissue, since this needed to be removed from the STOMP patterning device, and the swelling made this more difficult. The v-shaped degradable channel wall regions were patterned with 15 μl of hydrogel precursor each and filled until the liquid bulged slightly out of the channel. Devices with the patterned PEG in the v-shaped channels were incubated at 37°C for 30 minutes to allow for gelling. Sacrificial 1X PBS was used to prevent drying out of the hydrogel. After 30 minutes, devices were removed from the incubator and the cylindrical region of the patterning rails that surround the posts were treated with 1% BSA for one hour to prevent the tissue in contact with the circular walls from sticking upon removal of the patterning device. BSA was aspirated and devices were allowed time to dry for 5 to 10 minutes. Some absorption of the BSA by the hydrogel in the v-shaped degradable channel was observed, but swelling caused by this absorption resolved after 5 to 10 minutes of drying in open air. The center tissue region of the device was patterned with 50 μL of hydrogel precursor, then placed in the 37°C incubator to gel for 30 minutes. Sacrificial 1X PBS was added to the bottom of the well plates, beneath the patterning device, to prevent gels from drying out during gelation.

Once gelled, the hydrogel devices were inverted and completely submerged in 3 mL of eSrt(2A9) sortase solution. This sortase cleaves the peptide LAETG, which was used to pattern the degradable channel wall region in the v-shaped channels. The eSrt(2A9) sortase solution was prepared to a final concentration of 50 µM eSrt(2A9), 18 mM triglycine (GGG) (ChemImpex; Wood Dale, IL), and 1.8 mM CaCl2. GGG and CaCl2 were resuspended in cell culture media and adjusted to pH 7 using a 6 N solution of NaOH. Media was added to the final volume and the solution was sterile filtered with a 0.2 µm syringe filter. The devices were incubated in the eSrt(2A9) sortase working solution for 16 hours to degrade the PEG patterned in the v-shaped channel walls. Holes were added to the design to aid in permeation of the sortase solution into this region. After incubation, the clips and patterning device that now had a degraded channel wall were gently removed, releasing the PEG-based suspended hydrogel. Removal of the STOMP patterning device was performed while the hydrogel was submerged in a media. A video of the removal process can be seen in Supplementary Movie 9.

### EHT analysis

#### Force measurements

EHTs were placed in a Tyrode’s buffer (1.8 mM CaCl2, 1 mM MgCl2, 5.4 mM KCl, 140 mM NaCl, 0.33 mM NaH2PO4, 5 mM glucose, pH 7.35) at 37 °C for contractile analysis. Biphasic field stimulation at 1.5 Hz (5 V/cm for 10 ms duration) during imaging was provided by a custom pacing device incorporating carbon electrodes designed to fit in a 24-well plate and an electrical stimulator (Astro Med Grass Stimulator, Model S88X) as previously described[26,60,91]. Brightfield videos of EHT contraction were taken at 66.7 fps for 7.5 seconds for stimulated contractile measurements and 15 seconds for spontaneous contractile measurements. A custom MATLAB script was used to track the deflection of the flexible post and calculate the twitch force, shortening velocity, time to peak, time to 50% relaxation, time to 90% relaxation, beat frequency, and cross-sectional area. Specific force was calculated by dividing the twitch force of each tissue by its cross-sectional area.

#### Immunofluorescent Imaging

To visualize HS5-cells expressing GFP or mCherry which were seeded in the EHT outer or center regions respectively, EHTs were placed in a Tyrode’s buffer and imaged at 20 fps on an ORCA- Flash4.0 C13440 CMOS camera (Hamamatsu) on a Nikon TEi epifluorescent microscope with a FITC or TRITC filter cube. Images were processed using ImageJ.

### Periodontal tissue analysis

#### Calculation of PTC contractile force

Contractile force generated by PTCs was calculated by quantifying the magnitude of deflection they caused in the post pairs they were suspended between. Based on a modulus of elasticity of 2.5 MPa for PDMS, the bending stiffness (Kpost) of the flexible posts was calculated to be 0.95 μN/μm, as done previously[59]. Contractile force was calculated by multiplying the bending stiffness by the deflection of the post (initial length of tissue, L0, minus the final length of tissue, LF) (see Supplementary Figure 11).

#### Alizarin red staining

PTCs on posts were fixed in 4% PFA at room temperature for 1 hour and then removed from the posts. PTCs were then washed with DH2O, and incubated in alizarin red solution (pH 4.1 to 4.3) (Sigma-Aldrich) for 45 min at room temperature and washed 4x with DH2O on a shaker at room temperature. Whole PTCs were then coverslipped in VECTASHIELD mounting medium and allowed to dry overnight. The next day, brightfield microscope images at 4X were taken. Images were processed using the FIJI image analysis software.

#### Statistical analysis

Data from both the EHT and PTC work represents tissues collected from three independent experiments. All data points in the figures designate values for a single tissue. All values shown are reported as the mean ± standard error of mean. For the EHT work, results were compared by using an unpaired t-test with two tails. For the PTC work, results were compared using one-way ANOVA with post-hoc comparisons using GraphPad Prism. Differences with p-values ≤ 0.05 were considered statistically significant and denoted with an asterisk.

## Acknowledgements

This publication was supported by the National Institutes of Health (NIH) through R35GM128648 (ABT), R35GM138036 (CAD), R01HL149734 (NJS), R03DE029827 (TEP, NJS), T32CA080416 (IK), F30HL158030 (AJH), R90DE023059 (PM), 5TL1TR002318-08 (LGB), a diversity supplement to R35GM128648 (EEB), and the University of Washington. This work was partially supported by Friends of FSH Research and The Chris Carrino Foundation for FSHD (LGB) and fellowship funds from Senator Paul D. Wellstone Muscular Dystrophy Specialized Research Center - Seattle (NIAMS P50AR065139) (LGB). This work was also partially funded by a gift to support research from Ionis Pharmaceuticals (ABT, LGB, AJH). The content is solely the responsibility of the authors and does not necessarily represent the official views of the National Institutes of Health or other funders.

## Author Contributions

AJH, LGB, NJS, and ABT conceived the project. NJS and ABT supervised the project. AJH and LGB designed and fabricated the device with process inputs from JB, RCB, AW, EB, and ABT. AJG, PM and NAM fabricated the PDMS posts with process inputs from NJS. JB performed the theoretical analysis and derivations of the Laplace pressure at the pinning features. LGB and AJH performed the experimental pinning of collagen in the different pinning features. AJG designed and conducted the engineered heart tissue experiments with design inputs from LGB, AJH, and NJS. AJG performed the data analysis for the engineered heart tissue experiments. PM designed and PM and NAM conducted the periodontal tissue construct experiments with design inputs from AJH, LGB, TEP, and NJS. PM performed the fluorescent staining and data analysis of the periodontal tissue constructs with NAM and TPL. SHN designed the suspended core devices with design inputs from AJH. ARV designed the degradable wall devices with design inputs from AJH and IK. IK synthesized the degradable PEG components and assisted in experimental design with input from CAD. AJH, LGB, AJG, PM, JMW, AL, SHN, ARV, EEB, RMP, JCT and NAM assisted with cell culture. JMW and AL performed the contact angle measurements. AJH and LGB wrote the manuscript with significant inputs from AJG, PM, JB, SJT, CAD, TEP, EB, NJS, and ABT.

## Data and materials availability

All design files and all videos used to validate pinning theory (Figure 2 and Supplementary Figure 2) will be available publicly in figshare at https://doi.org/10.6084/m9.figshare.27905331, reference number 27905331. All other data are available in the main text or the supplementary materials.

