## Supplementary Information for "Suspended Tissue Open Microfluidic Patterning (STOMP)"

### Supporting Information for
#### Suspended Tissue Open Microfluidic Patterning (STOMP)

Amanda J. Haack[1,2]*, Lauren G. Brown[1]*, Alex J. Goldstein[3,4,5], Priti Mulimani[3,6], Jean Berthier[1], Asha R. Viswanathan[1], Irina Kopyeva[7], Jamison M. Whitten[1], Ariel Lin[1,8], Serena H. Nguyen[1], Thomas P. Leahy[3,9], Ella E. Bouker[1], Ruby M. Padgett[3,9], Natalie A. Mazzawi[3,6,10], Jodie C. Tokihiro[1], Ross C. Bretherton[3,7], Aaliyah Wu[1], Stephen J. Tapscott[11,12], Cole A. DeForest[1,3,7,8,13,14], Tracy E. Popowics[3,6], Erwin Berthier[1], Nathan J. Sniadecki[3,5,7,9 #], Ashleigh B. Theberge[1,15 #]

*Authors contributed equally to this work
#Corresponding Authors: Nathan J. Sniadecki & Ashleigh B. Theberge


**Affiliations:**

1. Department of Chemistry, University of Washington, Seattle, WA, 98195 USA
2. Medical Scientist Training Program, University of Washington School of Medicine, Seattle, WA, 98195 USA
3. Institute for Stem Cell & Regenerative Medicine, University of Washington, Seattle, WA, 98109 USA
4. Department of Materials Science and Engineering, University of Washington, Seattle, WA, 98195 USA.
5. Department of Laboratory Medicine & Pathology, University of Washington, Seattle, WA, 98195 USA
6. Oral Health Sciences, School of Dentistry, University of Washington, Seattle, WA, 98195 USA
7. Department of Bioengineering, University of Washington, Seattle, WA, 98195 USA
8. Molecular Engineering & Sciences Institute, University of Washington, Seattle, WA, 98109 USA
9. Department of Mechanical Engineering, University of Washington, Seattle, WA, 98195 USA
10. Department of Microbiology, University of Washington, Seattle, WA, 98195 USA
11. Human Biology Division, Fred Hutchinson Cancer Center, Seattle, WA 98109, USA
12. Department of Neurology, University of Washington, Seattle WA 98195, USA
13. Department of Chemical Engineering, University of Washington, Seattle, WA, 98195 USA
14. Institute for Protein Design, University of Washington, Seattle, WA, 98195 USA
15. Department of Urology, University of Washington School of Medicine, Seattle, WA, 98195 USA

**This PDF file includes:**

Supplementary Text
Figs. S1 to S13
Tables S1 to S3
Legends for Movies S1 to S9
Legends for Design Files S1 to S16
SI References (1-9)

**Other Supplementary Materials for this manuscript include the following:**

Movies S1 to S9
Design Files S1 to S16

**Supplementary Text**

Extended Technical Details of Methods

**1. Immunofluorescent imaging of periodontal tissues.** PTCs on posts were fixed in 4% PFA at room temperature for 1 hour. PTCs were then dismounted from the posts for the immunofluorescence staining process. In short, PTCs were permeabilized with 0.2% Triton-X for 10 min followed by blocking with 10% normal goat serum (Invitrogen) for 10 min at room temperature. Samples were then incubated for 1 hour with Alexa Fluor 647 phalloidin (Invitrogen A22287, 1:400), and Hoechst 33342 (ThermoFisher, 1:1000). After 1 hour, PTCs were rinsed three times in PBS for 10 min each on a room temperature shaker. Whole PTCs were mounted between glass slides and coverslips using VECTASHIELD mounting medium and allowed to dry overnight. A Leica SP8 confocal microscope was used for imaging. Three to four images per PTC were obtained under 10X objective at 2X zoom and 1024 x 1024 resolution. Since three independent experiments were carried out and each experiment was conducted in triplicate, we ensured a minimum of 30 images per condition for analysis. Laser strength and gain were kept constant between all samples and fields of view (Fig. S12).

**2. Synthesis of poly(ethylene glycol)-bicyclononyne (PEG-BCN).** 4-arm poly(ethylene glycol)-bicyclononyne (PEG-BCN) was synthesized as described in previous publications (*93-95*). 4-arm poly(ethylene glycol) tetraamine (MW 20kDa, 1.143 g, 0.0571 mmol, 0.2 mol $NH_2$ groups, 1x; JenKem Technology USA; Plano, TX), (1R,8S,9s)-bicyclo[6.1.0]non-4-yn-9-ylmethyl (2,5-dioxopyrrolidin-1-yl) carbonate (BCN-OSu) (Sigma Aldrich; St. Louis, MO) (100 mg, 0.343 mmol, 1.5x to $NH_2$ groups), and N,N-Diisopropylethylamine (DIEA, 159µL, 8 mmol, 4x to $NH_2$ groups) were dissolved in anhydrous dimethylformamide (DMF, 5 mL) and stirred overnight. The next day, the mixture was diluted with DI $H_2O$ (10x volume) and dialyzed overnight in DI $H_2O$ (molecular weight cutoff ~ 2kDa; SpectraPor, Repligen; Waltham, MA) and lyophilized over three days to yield a white powder. The powder was resuspended in sterile PBS at a 10 mM stock concentration and stored at -80°C until further use. $^1H$ NMR confirmed functionalization to be >95% by comparing integral values for characteristic BCN peaks (δ 2.24, 1.57, 1.34, 0.92) with those from the PEG backbone (δ 3.63).

**3. Peptide Synthesis of eSrtA(4S9)- and eSrtA(2A9)-sensitive Crosslinkers.** The eSrtA(4S9)- and eSrtA(2A9)-sensitive diazide peptide crosslinkers H-RGPQGIWGQLPESGGRK(dde)-$NH_2$ and H-RGPQGIWGQLAETGGRK(dde)-$NH_2$, respectively, were synthesized on rink amide ProTide resin (CEM Corporation; Charlotte, NC) following induction heating-assisted Fmoc solid-phase techniques with HCTU activation (Gyros Protein Technologies PurePep Chorus; Tucson, AZ) at a 0.4 mmol scale. Deprotection of the 1-(4,4-dimethyl-2,6-dioxacyclohexylidene)ethyl (dde) group was accomplished by treating with 2% hydrazine monohydrate in DMF (3x10 min). For azide modification of the both the N-terminal amine and the ε-amino group of the C-terminal lysine, 4-azidobutanoic acid (227 µL, 2 mmol, 4x to $NH_2$ groups), hexafluorophosphate azabenzotriazole tetramethyl uronium (HATU, 750 mg, 1.97 mmol, 3.95x to $NH_2$ groups) and DIEA (1.38 mL, 8 mmol, 16x to $NH_2$ groups) were prereacted for 5 minutes and then reacted with the peptide for 1.5 hours. For peptide cleavage and deprotection, the resin was treated with trifluoracetic acid/triisopropylsilane/water (95:2.5:2.5) for 3 hours, then crashed out and washed in ice-cold diethyl ether (2 x 150mL). The crude peptides were purified via semi-preparative reversed-phase high performance liquid chromatography with a linear gradient of 5-100% acetonitrile and 0.1% TFA for 45 minutes and then lyophilized to yield white powders of the final peptides $N_3$-RGPQGIWGQLPESGGRK($N_3$)-$NH_2$ and $N_3$-RGPQGIWGQLAETGGRK($N_3$)-$NH_2$. Peptide mass was verified via ESI-LCMS.

**4. eSrtA(4S9) and eSrtA(2A9) Expression and Purification.** The pET29B expression plasmids for eSrtA(4S9) and eSrtA(2A9) were a generous gift from Dr. David Liu at Harvard University (Addgene plasmids #75146 and #75145). The sortase variants were expressed and purified as previously described (*93*). Electrically competent BL21 cells were transfected with the eSrtA(4S9) and eSrtA(2A9) plasmids and selected on kanamycin-containing agar plates. 5 mL of Luria Broth (LB) with kanamycin (50µg/mL) was inoculated with a plasmid-containing colony and grown overnight at 37ºC shaking at 200 rpm. The following day, 1 L of LB broth, including 40 mL of autoinduction sugars (60% v/v glycerol, 10% w/v glucose, 8% w/v lactose) as well as kanamycin, was inoculated with the 5mL overnight culture and allowed to incubate at 37ºC, 200 rpm overnight. The following day, cells were pelleted via centrifugation (4000g, 20 mins), resuspended in lysis buffer (20 mM Tris, 50 mM NaCl, 10 mM imidazole; pH 7.5) with 1 mM phenylmethylsulfonyl (PMSF; TCI, Portland, OR) protease inhibitor, and lysed by sonication (6x, 3 min cycle, 30% amplitude; Fisher Scientific; Waltham, MA). The lysate was clarified via centrifugation (20 mins, 11000g). The clarified lysate was then purified using an ÄKTA Pure 25 L FPLC (Cytiva; Marlborough, MA) equipped with a 5 mL HisTrap HP column at a flow rate of 5 mL $min^{-1}$. The column was equilibrated with 5 column volumes (CV) of lysis buffer, followed by loading the sample and washing with 8 CV of endotoxin removal buffer (20 mM Tris, 50 mM NaCl, 20 mM imidazole, 0.1% Triton-X 114; pH = 7.5) and 8 CV of wash buffer (20 mM Tris, 50 mM NaCl, 20 mM imidazole; pH = 7.5). His-tagged protein was eluted over an 8 CV gradient of imidazole (20 mM Tris, 50 mM NaCl, 5-250 mM imidazole; pH = 7.5) into a 96 well plate; protein-containing fractions were pooled and dialyzed with SnakeSkin dialysis tubing (10 kDa cut off; ThermoFisher, Waltham, MA) in PBS over two days. Purified sortase was spin-concentrated using an Amicon Ultra 15 centrifugation filter (10,000 Da cut off; Sigma Aldrich; St. Louis, MO). The final stocks were diluted with PBS containing 10% glycerol to a concentration of 100 µM. Sortase purity was evaluated with sodium-dodecyl sulfate-polyacrylamide gel electrophoresis and identity was confirmed with electrospray ionization mass spectrometry on an AB SCIEX 5600 QTOF instrument (SCIEX; Framingham, MA).

Extended Technical Descriptions of Results

**1. Full details on theoretical characterization of pinning in STOMP channels.**

In this study, we investigate the successive pinning of the two liquids in the STOMP device shown in Fig. S3. The “green” liquid represents a hydrogel (e.g., cell-ECM mixture) first pipetted into the outer cylindrical regions of the STOMP device. The “orange” liquid represents a second hydrogel (cell-ECM mixture) pipetted later in the center region. In this analysis, we assume both hydrogels (green and orange regions) are 5 mg/mL collagen, but this analysis can be extended to any hydrogel by utilizing that specific hydrogel’s properties (e.g., density, contact angle on patterning device surface, etc.).

It is assumed that liquid #1 (green in Fig. S3) is pipetted first, so it is the pinning of this liquid that is investigated here. In a first approach, we follow a 2D approach, neglecting the effect of the horizontal free surfaces at bottom and top. We shall justify this approximation later in the text. Here, we analyze two pinning feature designs we refer to as convex “vampire” pins and concave “cavity” pins.

We first address the case of the convex “vampire” pinning features (Fig. S4). The design is constituted by two facing reliefs resembling two “teeth”. We note that $\alpha$ is the angle formed by the triangular-shaped relief, $\theta$ the contact angle of the hydrogel with the channel wall, and w is the distance between the two facing reliefs of the convex “vampire” pins. The Laplace pressure $\Delta P$ of the liquid is given by equation (S1), where r is the curvature radius of the pinned interface (in the horizontal plane) and denotes the surface tension of the liquid.

$$\Delta P = \frac{\gamma}{r} \qquad \text{S1}$$

At the pinning limit (i.e., the maximum interface the fluid bulges before pinning is broken), the bulging fluid front forms the angle $\theta$ with the external side of the triangular edge (*96-100*); its complement is noted in Fig. S4. Any additional bulging results in a capillary flow passing over the pinning feature. Let us denote the points A and B the tip of the “teeth”, O the center of the circular arc formed by the interface, and C the middle of the segment AB. Using the construction in Fig. S4, the angle {AC,AO} is simply given by the expression $\pi/2-\Theta+\alpha/2$. Then considering the rectangular triangle ACO, we find

$$\cos\left(\frac{\pi}{2} - \theta + \frac{\alpha}{2}\right) = \frac{w}{2r} \qquad \text{S2}$$

or

$$\sin\left(\theta - \frac{\alpha}{2}\right) = \frac{w}{2r} \qquad \text{S3}$$

Remark that we have considered ϴ>α/2. When ϴ=α/2, the fluid interface is flat, and r is infinite. Substituting equation (S3) into equation (S1) yields the threshold Laplace pinning pressure, above which pinning is lost in the convex "vampire" pinning feature design (Eq. S4).

$$\Delta P = \frac{\gamma}{r} = \frac{2\gamma \sin\left(\theta - \frac{\alpha}{2}\right)}{w} \qquad \text{S4}$$

Second, we analyze the case of the concave "cavity" pinning features (Fig. S5). The Laplace pressure is still given by equation (1). Let us note β as the angle of solid at the ridge. Let us separate two cases: β > θ and β < θ. The first case is the usual case, because θ is generally small and the cavity angle β cannot be too small due to the 3D printing fabrication process of the STOMP device. The first case (β > θ) corresponds to the configuration used here in this work.

<u>Case β > θ:</u> Again, at the limit, the bulging angle forms the angle θ with the external side of the tooth4–8; its complement is noted as π-θ in Fig. S5. Any additional bulging results in a capillary flow passing over the ridge. Let us denote the points A and B the tip of the ridges, O the center of the circular arc formed by the interface, and C the middle of the segment AB. Using the construction in Fig. S5, the angle {AC,AO} is simply β-θ. Then considering the rectangular triangle ACO, we find

$$\cos(\theta - \beta) = \frac{w}{2r} \qquad \text{S5}$$

Substituting equation (S5) into (S1) yields the threshold Laplace pinning pressure, above which pinning is lost in the concave "cavity" pinning feature design (Eq. S6).

$$\Delta P = \frac{\gamma}{r} = \frac{2\gamma \cos(\theta - \beta)}{w} \qquad \text{S6}$$

<u>Case β < θ:</u> While this case is not the usual case, we mention it for completion of the study. In this case, the fluid front creates a bulging curvature, where the center of the curvature (point O in Fig. S6) formed is located past the pinning ridge (line segment AB in Fig. S6). The threshold Laplace pinning pressure is still the same as Eq. S5, but this equation reaches its maximum value when β is equal to θ, which is 30° in our theoretical model (the average contact angle of 5 mg/mL collagen on 3D printed resin treated with 1% BSA). Therefore, the maximum Laplace pinning pressure that can be experienced in the concave "cavity" pinning feature is given below in equation (S7) and is visualized as the solid green line in Fig. S6.

$$\Delta P = \frac{2\gamma}{w} \qquad \text{S7}$$

Lastly, because gravity has a small, but not negligible effect, we must also consider the hydrostatic pressure at the pinning interface. For both pinning feature designs, the hydrostatic pressure at the bottom of the channel is given below in equation (S8), where $\rho$ is the density of the hydrogel, g is the gravitational constant, and h is the height of the channel.

$$\Delta P = \rho g h \qquad \text{S8}$$

**2. Analyzing the 3D effect of gravity in theoretical approach to pinning in STOMP**

In our theoretical pinning model for STOMP, we used a 2D approach where we neglected the effect of the horizontal free air-liquid interfaces at the bottom and top of the open channel. To justify this approach, we also analyzed the potential 3D effects, which are twofold: 1) gravity effects that may distort the bottom air-liquid interface of the open channel and 2) corner effects from the rounded vertical air-liquid interface along the height of the open channel.

First, to analyze the effect of gravity, the Bond number has to be calculated. For testing the pinning in STOMP, devices with a channel height of 3.5 mm and width of 1.2 mm were used. The equation for bond number Bo is given below in equation (S9), where $\rho$ is the density of the 5 mg/mL collagen hydrogel, g is the gravitational constant, h is the height of the channel, R is the radius of the outer cylinders, and $\gamma$ is the surface tension of the liquid (estimated to be water here).

$$Bo = \frac{\rho g h R}{\gamma} \qquad \text{S9}$$

In the case of the outer cylinders, the radius is 1.5 mm. Therefore, the bond number for the outer cylinders can be approximately calculated, as shown below in equation (S10).

$$Bo_{outer\ cylinders} = \frac{958\ \frac{kg}{m^3} \times 10 \frac{m}{s^2} \times 3.5 \cdot 10^{-3} m \times 1.5 \cdot 10^{-3} m}{72 \cdot 10^{-3} \frac{N}{m}} \approx 0.7 \qquad \text{S10}$$

In the case of the central region of the open channel, the radius R is replaced by the semi-width, w/2. When experimentally testing the pinning, we used the STOMP devices with a channel height of 3.5 mm and a width of 1.2 mm. Therefore, the bond number for the central region can be approximately calculated with a semi-width of 0.6 mm, as shown below in equation (S11).

$$Bo_{central\ region} = \frac{958 \frac{kg}{m^3} \times 10 \frac{m}{s^2} \times 3.5 \cdot 10^{-3} m \times 0.6 \cdot 10^{-3} m}{72 \cdot 10^{-3} \frac{N}{m}} \approx 0.3 \qquad \text{S11}$$

In both cases of the outer cylinders that surround the posts and the central region of the open channel where flow occurs, we calculated that Bo < 1, which means surface tension forces dominates in the open channel. In both cases, gravity has a small effect on the horizontal free air-liquid interface at the bottom and top of the open channel. While the effect of gravity is not negligible, it is small enough that we can justify using a 2D approach in our theoretical characterization of the pinning in the STOMP devices.

The corner effects of the vertical air-liquid interface along the height of the open channel are more difficult to predict. We used Brakke's software Surface Evolver to determine the shape of these interfaces (*101*). The results are plotted in Fig. S7, which demonstrates that some corner effects do exist in our STOMP system.

**3. Experimentally testing different volumes and pinning feature angles in STOMP**

To understand the sensitivity of our system to changes in volume, we tested pinning in five different pinning angles of both pinning feature designs (i.e., α=10°, 20°, 30°, 45°, and 60° for the convex "vampire" pins and $\beta$=40°, 60°, 90°, 100°, and 120° for the concave "cavity" pins, n=3 devices for each pin angle) at four different volumes (20, 22, 23, and 24 μL). These results are illustrated in Fig. S2.

Every volume tested exhibited different pinning results, where increasing the volume resulted in less successful pinning. At every volume, the concave "cavity" pinning feature with pins $\beta$=120° resulted in failed pinning; this matches what we would expect from the theoretical results (see Fig. 2d). Generally, in all volumes tested, the concave "cavity" pinning feature was more consistent than the convex "vampire" pinning feature. This phenomenon may be explained by the large range in the pinning Laplace pressure for the convex pinning features (see Fig. 2c). We observed that a volume of 23 μL best fitted our theoretical model.

When 23 μL was pipetted into the convex "vampire" pinning features, we observed successful pinning of the collagen solution when α=10°, 20°, 30° (n=3/3 devices pinned). These experimental results match what would be expected from the average theoretical maximum Laplace pinning pressure (Fig. 2c). Further, there was less successful pinning when α=45° and 60° (n=2/3 devices did not pin). We note that one out of the three devices for both the angles α=45° and 60° had successful pinning, but these angles have conditions of success and failure within the pinning range of the maximum Laplace pinning pressure (Fig. 2c, green shading showing successful conditions and magenta shading showing failure conditions), suggesting that those devices had variable BSA adsorption onto the channel walls that may have changed the contact angle of the collagen on the channel surface.

When 23 μL was pipetted into the concave "cavity" pinning features, we observed successful pinning of the collagen solution when $\beta$=40° and 60° (n=3/3 devices pinned) and no pinning when $\beta$=90°, 100°, and 120° (n=3/3 devices did not pin). While successful pinning at $\beta$=40° and 60° experimentally matches the theory, loss of pinning was observed in all three replicates in devices with $\beta$=90° which was expected, even within the contact angle range, to successfully pin (Fig. 2d). It is possible that the contact angle was towards the lower bound of the possible threshold Laplace pressure (Fig. 2d), where the line does cross below the hydrostatic pressure at $\beta$=90°. There could also be additional reasons for fluid de-pinning when not predicted. In some cases, a fluid may initially pin, but then lose that pinning if there are deformities present on the pinning feature (i.e., rough surface, inconsistency in 3D printing, etc.).

**4. Influence of liquid volume in pinning**

Since we observed differences in pinning as the pipetted volume changed, we wanted to look at two possible factors that could influence pinning: 1) change in the horizontal interface of the liquid in the channel (i.e., change in volume of liquid bulging at the top and bottom of the channel) and 2) change in the vertical interface of the pinned fluid front (i.e., change in the pinned fluid angle).

First, we will estimate the volume bulging at the bottom of the channel due to the weight of the precursor liquid hydrogel in the STOMP channel. At the bottom of the channel, the air-liquid interface bends due to hydrostatic pressure, as depicted in Fig. S8. If we consider the bending of the horizontal bottom surface to be a cylinder, then the curvature of the surface is $\kappa$ = 1/R. Therefore, the bending of the surface by hydrostatic pressure can be modeled by equation (S12), where R is the curvature radius of the bulging surface.

$$\Delta P = \rho g h = \frac{2\gamma}{R} \quad \text{S12}$$

If $a$ denotes the radius of the cylinder, the volume of the liquid in the spherical cap bulging below the cylinder is given below in equation S13 (*96*).

$$V_{horizontal\ bulging} = \frac{\pi}{6}\left(R - \sqrt{R^2 - a^2}\right)\left[3a^2 + \left(R - \sqrt{R^2 - a^2}\right)^2\right] \quad \text{S13}$$

From Berthier and Brakke (*96*), the negative bulging height can be deduced as

$$R = \frac{a^2 + h^2}{2h} \quad \text{S14}$$

or

$$h = R - \sqrt{R^2 - a^2} \quad \text{S15}$$

If $\rho$ is 958 kg/m3 (density of 5 mg/mL collagen), g is 9.8 m/s$^2$, h is 3.5 mm (the height of the open channel), $\gamma$ is 72 mN/m, and a is 1.5 mm (the radius of the outer cylinders of the STOMP device), then we find that the curvature radius R is 4.38 mm, giving a bulging volume of 0.945 µL (or 0.945 mm$^3$) with a bulging height of 0.265 mm (or 265 µm). Thus, if the volume of the precursor hydrogel is equal to exactly the volume of the cylinder plus the inner channel walls (unto the pinning features), then the bottom surface bulges out 265 µm below the channel.

Next, we will calculate the surface area of the pinned fluid's vertical interface as a function of the pinning angle. Figure S9 shows a top-down cross section of the STOMP channel, where R is the curvature radius of the bulging vertical interface, w is the width of the open channel, $\delta$ is the bulging distance of the pinned fluid's vertical interface, $\phi$ is the pinning angle of the fluid front, and S is the vertical surface denoted with a dark green line. In a 2D approximation, a change in volume delimited by the surface S can be modeled as

$$V_{vertical\ bulging} = h \times S \quad \text{S16}$$

where h is the height of the open channel. Geometrical considerations give us the curvature radius R, denoted below in equation (S17).

$$R = \frac{w}{2\sin(\phi)} \quad \text{S17}$$

Modeling the vertical bulging surface as a circular segment, the surface area of the arc (denoted as S) can be modeled by equation (S18).

$$S = \frac{R^2}{2}\left[2\phi - \sin(2\phi)\right] \quad \text{S18}$$

Substituting equation (S17) and (S18) into (S16), we obtain an equation that calculates the change in volume of the vertical surface S as a function of the bulging pinning angle $\phi$.

$$V_{vertical\ bulging} = h \times \frac{R^2}{2}\left[2\phi - \sin(2\phi)\right] = h \times \frac{w^2}{8} \times \frac{2\phi - \sin(2\phi)}{\sin^2(\phi)} \quad \text{S19}$$

Additionally, the bulging distance $\delta$ is given by equation (S20).

$$\delta = R\left[1 - \cos(\phi)\right] = \frac{w(1 - \cos(\phi))}{2\sin(\phi)} \quad \text{S20}$$

As more liquid is added to the channel, the vertical interface of the pinned fluid will continue to bulge past the pinning features, thus changing the angle of the pinned liquid. Let us now assume that an additional liquid volume, denoted as dV, is added to the initial volume V (Fig. S10). The surface area of the vertical interface now becomes S + dS. To find the effect of the increase of volume dV on the bulging angle $\phi$, we can calculate the derivative of equation (S19) to find dV/d$\phi$. This derivatization gives us equation (S21)

$$\frac{dV}{d\phi} = h \times \frac{w^2}{4} \times \frac{[\sin(\phi) + 2\phi + \cos(3\phi)]}{\sin^3(\phi)} \qquad \text{S21}$$

which gives us $d\phi/dV$ in equation (S22).

$$\frac{d\phi}{dV} = \frac{4\sin^3(\phi)}{h \times w^2 \times [\sin(\phi) + 2\phi + \cos(3\phi)]} \qquad \text{S22}$$

The values of the increased bulging angle $\phi$ with an increase of liquid volume of only 1 µL (or 1 $mm^3$) for 4 initial pinning angles $\phi_0$ (20°, 30°, 40°, and 50°) are listed in Table S2. We find that adding 1 µL of additional volume can change the bulging angle; for example, an initial pinning angle of 40° or 50° can increase by 17° or 28°, respectively, with the addition of only 1 µL. This calculation can help explain why experimentally we observed changes in successful pinning of the fluid front at different pipetted volumes.

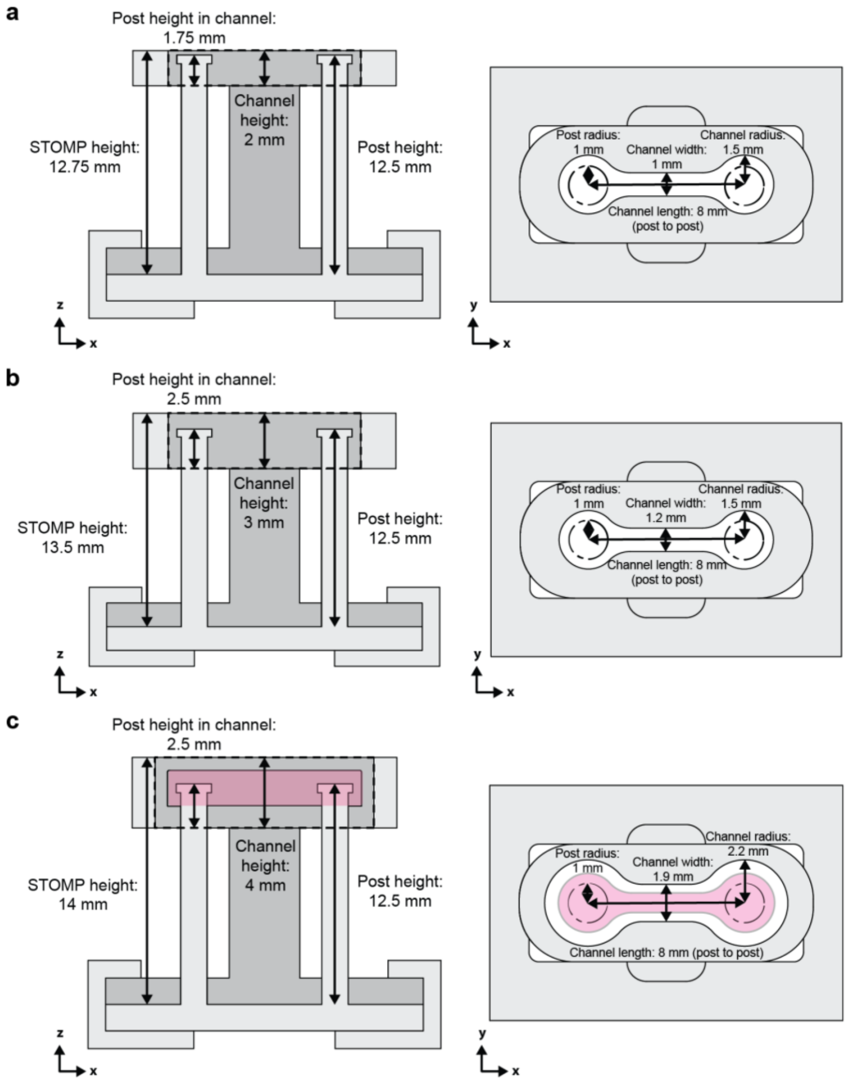


**Fig. S1.** Dimensions of STOMP devices used in this work. a) 2 mm height device, b) 3.5 mm height device, and c) outer region patterning device surrounding previously patterned hydrogel (in pink), featured in Fig. 5.

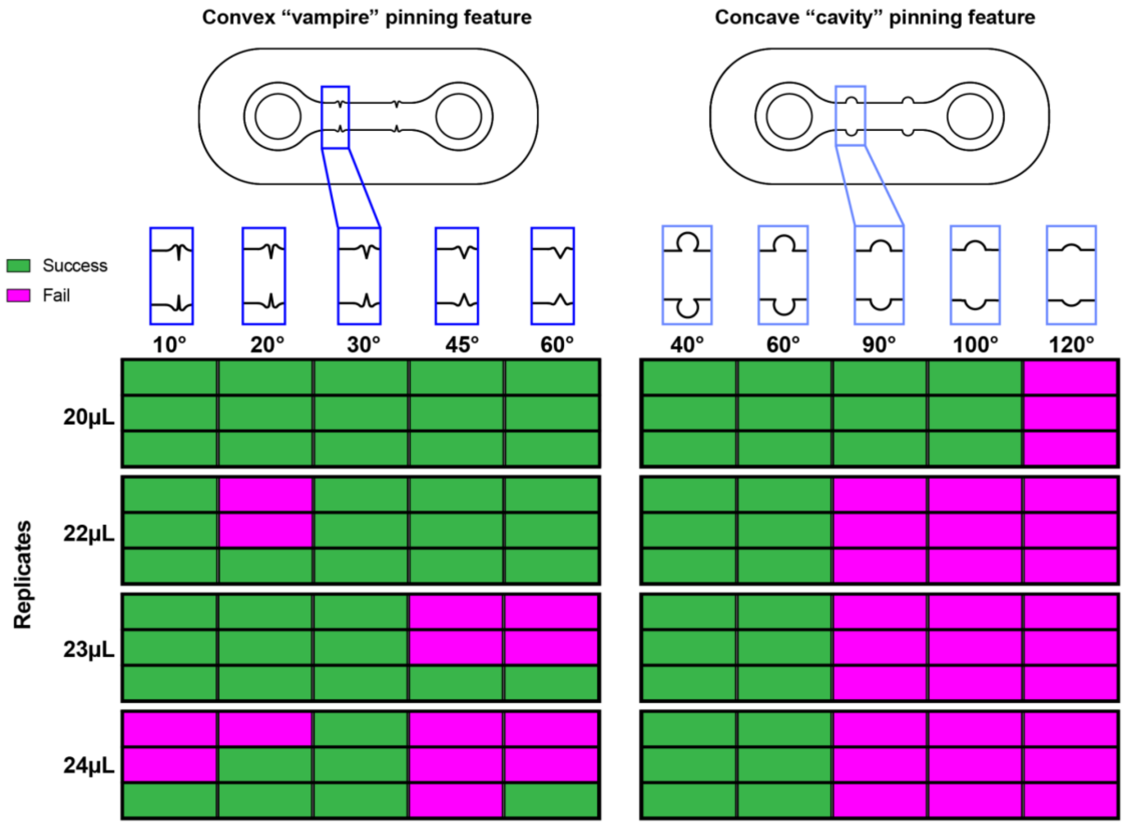


**Fig. S2.** Experimental results of pinning of the 5 mg/mL collagen precursor fluid front at different pipetted volumes. For both the convex "vampire" and concave "cavity" pinning features, five different pinning feature angles were tested to observe either success (green) or failure (magenta) of pinning at the pinning feature. For the convex "vampire" pinning features, the angle of the pinning feature refers to the vertex angle of the triangular pinning feature. For the concave "cavity" pinning features, the angle of pinning feature refers to the angle between the line tangent to the semicircular pinning feature to the straight channel wall.

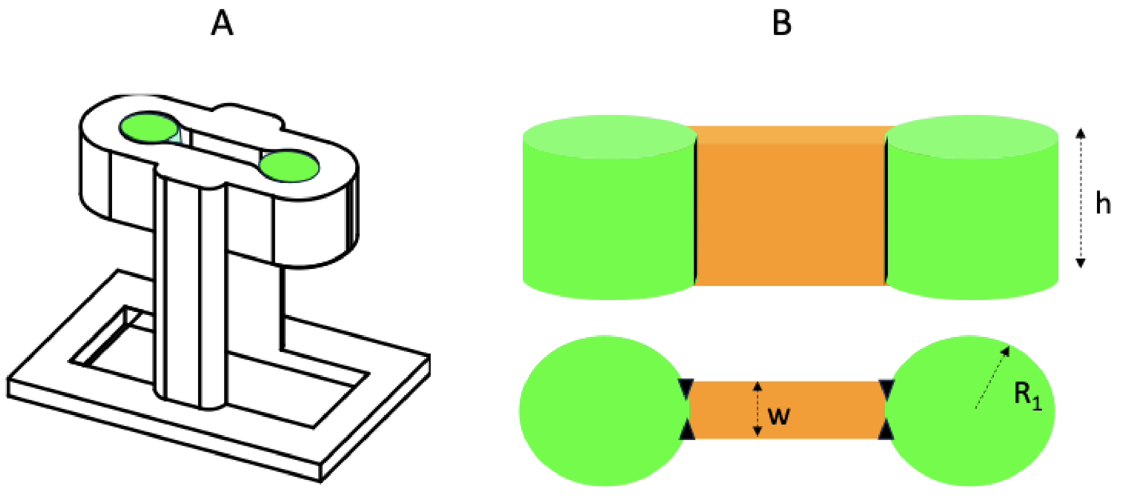


**Fig. S3.** Modeling fluid in the multi region STOMP device. A) isometric view of the device. B) top: side view of the filled device; bottom: top view of the filled device. The black lines and arrows symbolize the relief/design that pins the liquids during the filling.

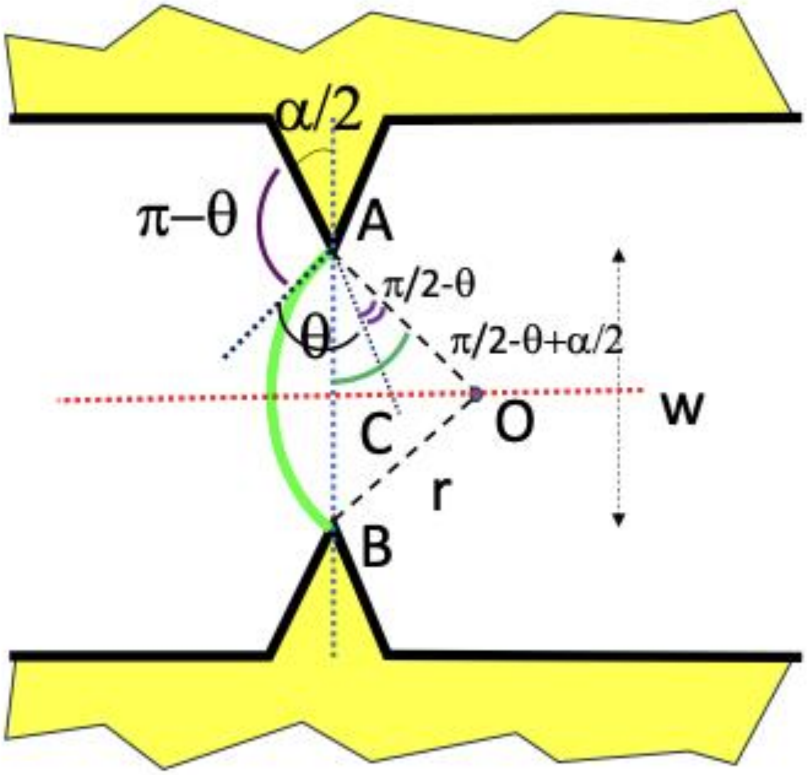


**Fig. S4.** Geometrical analysis of convex "vampire" pinning features. Green line is the fluid front, where the hydrogel precursor was pipetted first into the region on the right, so the fluid front is bulging to the left at the Laplace pinning threshold.

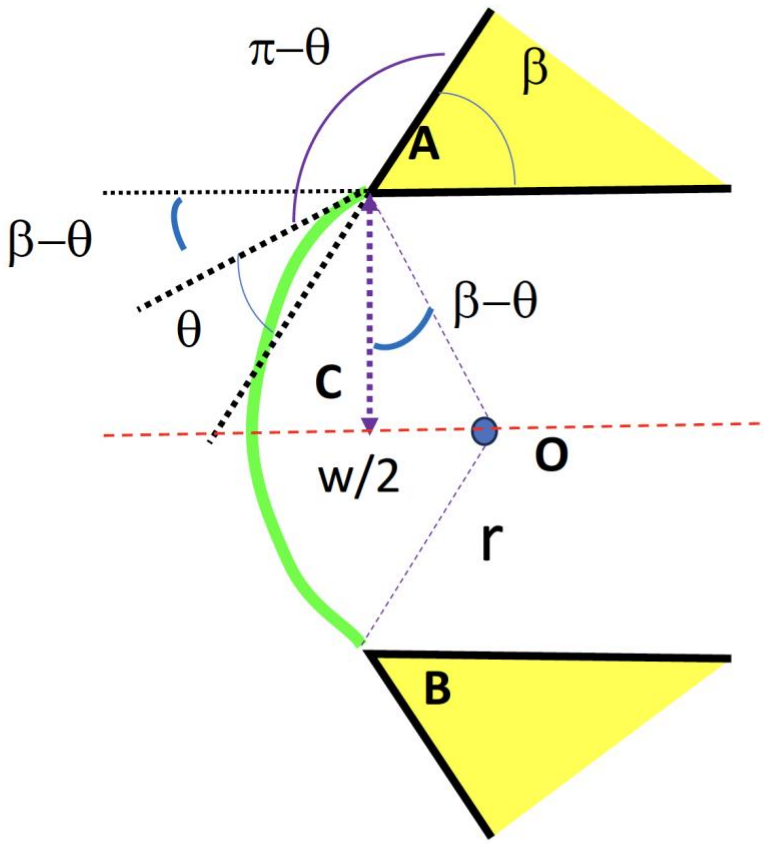


**Fig. S5.** Geometrical analysis of concave "cavity" pinning features when β > θ. Green line is the fluid front, where the hydrogel precursor was pipetted first into the region on the right, so the fluid front is bulging to the left at the Laplace pinning threshold.

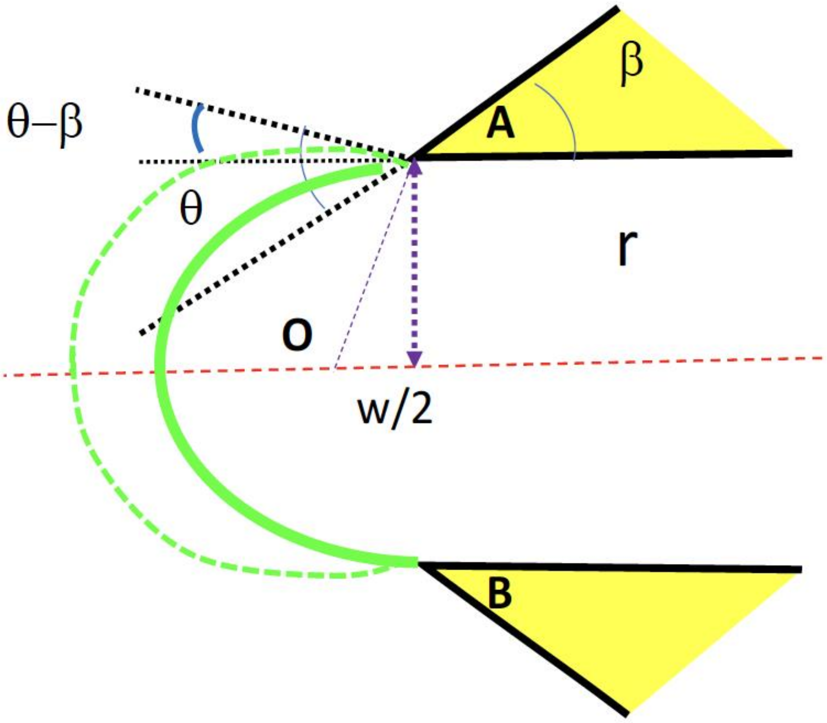


**Fig. S6.** Geometrical analysis of concave "cavity" pinning features when $\beta < \theta$. The dotted green line is the expected fluid front, where the hydrogel precursor was pipetted first into the region on the right, so the fluid front is bulging to the left. However, the actual fluid front is the solid green line, which is the maximum Laplace pinning threshold.

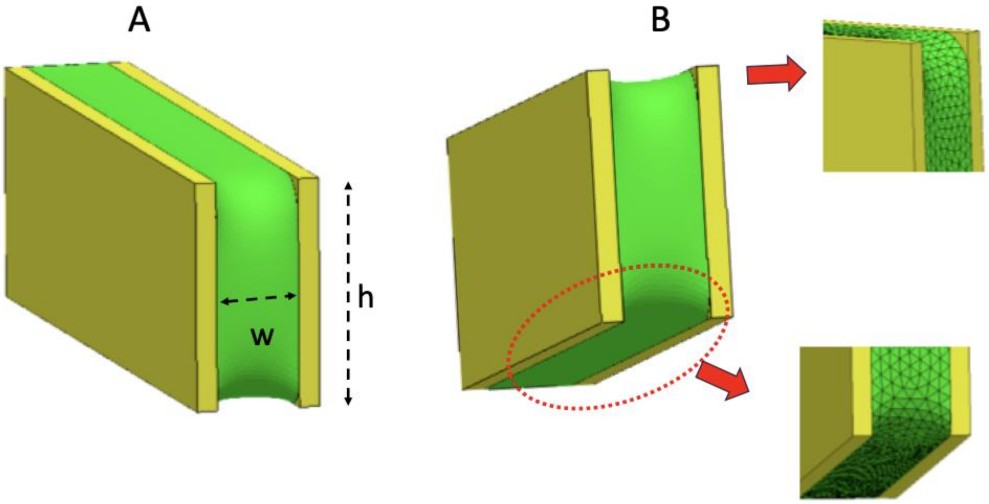


**Fig. S7.** 3D calculation of the free air-liquid interface of a liquid between two vertical walls with a width of 1.2 mm and a height of 3.5 mm. A) Isometric view of a fluid (green) within an open channel with two walls and no ceiling or floor. B) Closer view of the top and bottom corners of the vertical interface, showing the plots obtained using the Surface Evolver software.

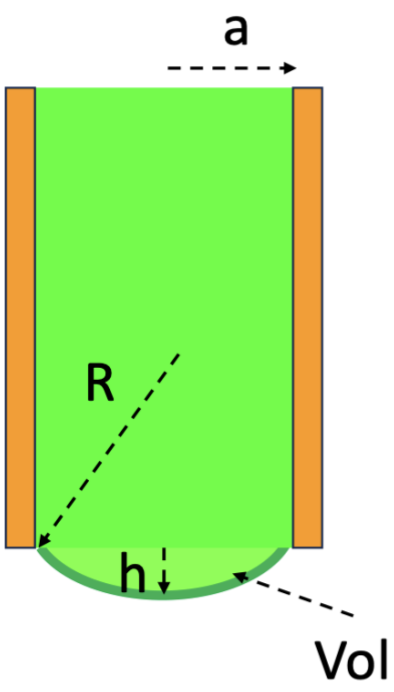


**Fig. S8.** Schematic of the fluid (green) bulging below the suspended channel walls (orange).

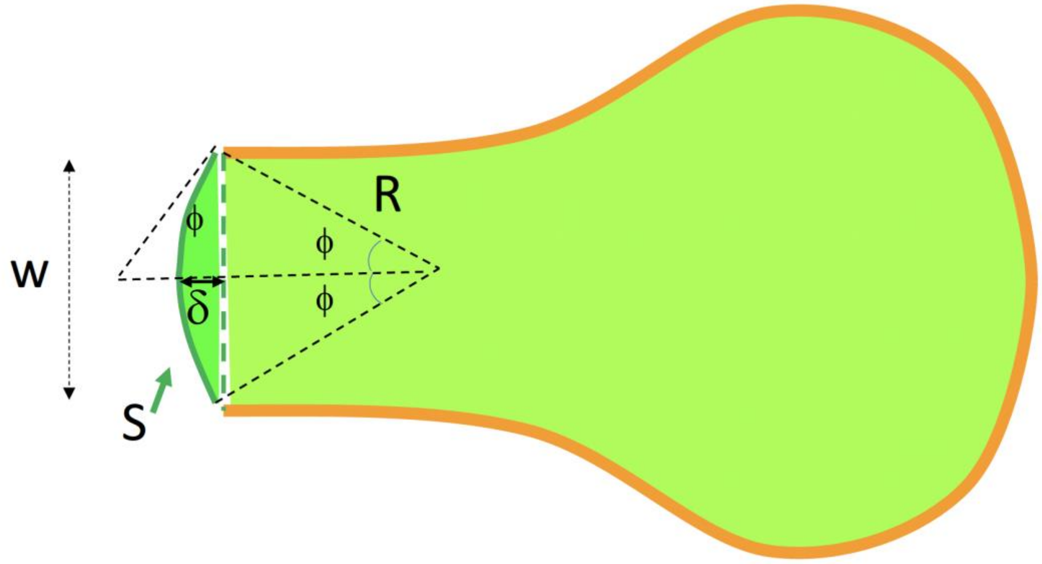


**Fig. S9.** Top-down schematic of the fluid (green) filling the outer regions of the suspended channel (walls denoted in orange) before reaching the pinning features (location marked by dotted green line). After reaching the pinning feature, the fluid will bulge some distance δ along the vertical interface, with the bulging angle denoted as $\phi$. The surface area S (dark green) is the surface area of the bulging liquid along the vertical interface.

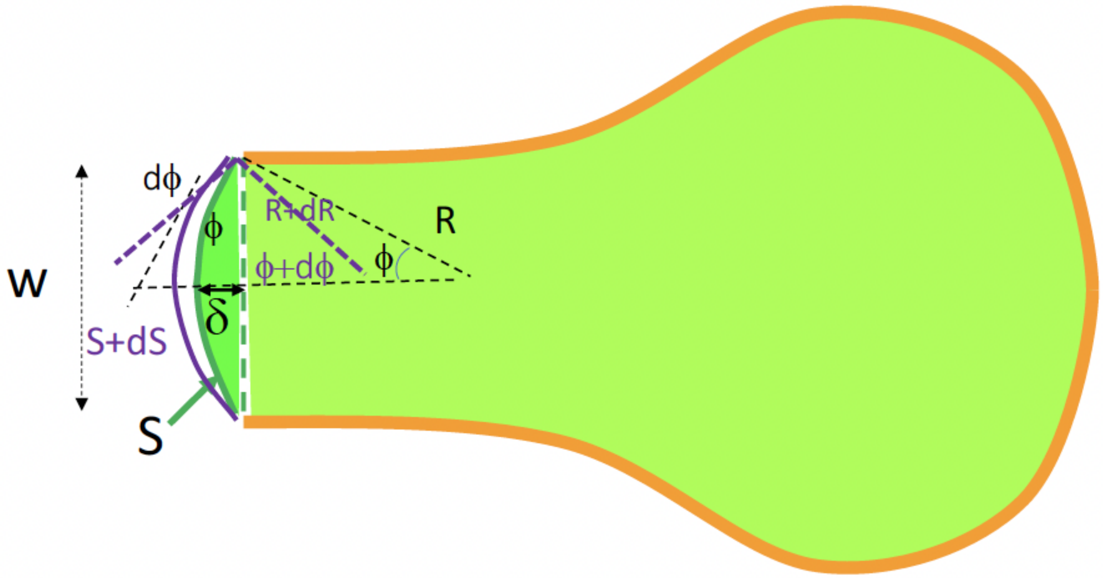


**Fig. S10.** Top-down schematic of how the bulging fluid (dark green) along the vertical interface changes after pipetting additional volume into the suspended channel.

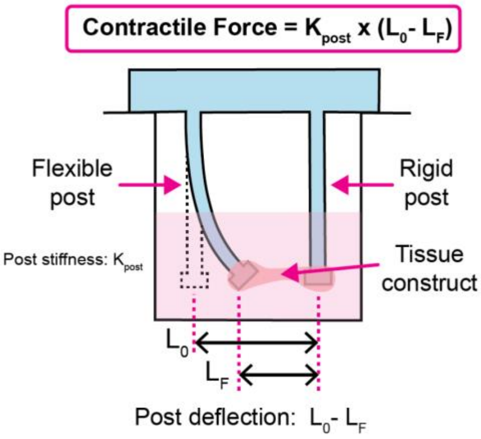


**Fig. S11.** Tissue contractile force calculation using post deflection. Deflection of post is measured by initial length of tissue, $L_0$, minus the final length of tissue, $L_F$.

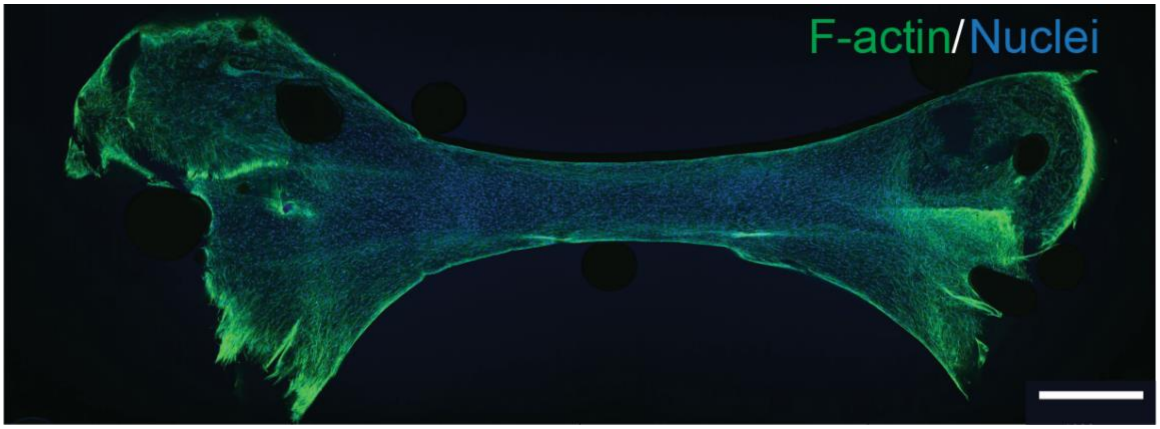


**Fig. S12.** Fluorescent image of a patterned PTC stained for F-actin (green) and nuclei (blue). Scale bar is 1 mm.

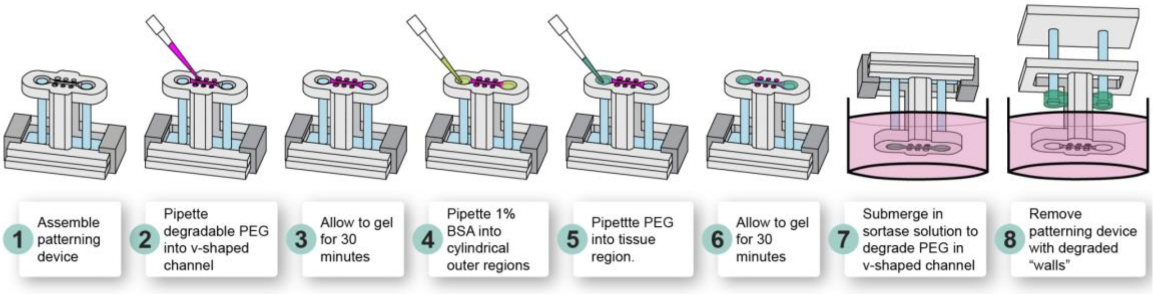


**Fig. S13.** Schematic showing process for patterning a degradable hydrogel channel wall and middle tissue region. The hydrogel depicted in magenta uses a sortase degradable crosslinker allowing for selective degradation of the channel wall with the addition of sortase.

Table S1. Experimental contact angle measurements of 5 mg/mL collagen on 1% BSA treated 3D printed resin.

| **Experiment number** | **1% BSA treated 3D printed surface number** | **Number of technical measured replicates from Krüss drop shape analyzer goniometer** | **Mean contact angle (degrees) from Krüss drop shape analyzer goniometer** | **Standard deviation (degrees) of contact angle from Krüss drop shape analyzer goniometer** | **Overall Average of mean contact angles (degrees)** | **Average standard deviation of contact angle (degrees)** |
|---|---|---|---|---|---|---|
| 1 | 1 | 10 | 34.73 | 0.98 | 29.62 | 3.51 |
| | 2 | 6 | 28.69 | 4.13 | | |
| | 3 | 9 | 48.41 | 2.07 | | |
| 2 | 4 | 6 | 18.11 | 2.55 | | |
| | 5 | 5 | 26.01 | 2.7 | | |
| | 6 | 10 | 36.31 | 2.46 | | |
| | 7 | 7 | 23.18 | 2.15 | | |
| | 8 | 5 | 20.82 | 1.36 | | |
| | 9 | 9 | 22.38 | 2.77 | | |
| 3 | 10 | 5 | 19.80 | 11.96 | | |
| | 11 | 7 | 41.85 | 1.93 | | |
| | 12 | 7 | 30.75 | 2.26 | | |
| | 13 | 8 | 39.64 | 3.6 | | |
| | 14 | 9 | 35.74 | 4.04 | | |
| | 15 | 5 | 21.75 | 2.62 | | |
| | 16 | 9 | 32.36 | 2.3 | | |
| | 17 | 5 | 39.81 | 2.81 | | |
| | 18 | 7 | 28.28 | 2.98 | | |
| | 19 | 6 | 29.06 | 4.45 | | |
| | 20 | 5 | 14.62 | 2.93 | | |

**Table S2. Change in the bulging angle of the pinned fluid's vertical interface as a function of change in volume.**

| w (mm) | h (mm) | $\phi_0$ (°) | $\phi_0$ (rad) | $\delta_0$ (μm) | $S_0$ ($mm^2$) | $V_0$ ($mm^3$) | $dV/d\phi$ ($mm^3$/rad) | $d\phi/dV$ (°/$mm^3$) | dV ($mm^3$) | $\phi$ (°) | $\delta$ (μm) |
|---|---|---|---|---|---|---|---|---|---|---|---|
| 0.8 | 3.5 | 20 | 0.35 | 70.5 | 0.04 | 0.13 | 21.56 | 2.66 | 1.0 | 23 | 80.1 |
| 0.8 | 3.5 | 30 | 0.52 | 107.2 | 0.06 | 0.20 | 6.93 | 8.27 | 1.0 | 38 | 138.8 |
| 0.8 | 3.5 | 40 | 0.70 | 145.6 | 0.08 | 0.28 | 3.25 | 17.66 | 1.0 | 57 | 220.2 |
| 0.8 | 3.5 | 50 | 0.87 | 186.5 | 0.10 | 0.36 | 2.05 | 27.95 | 1.0 | 78 | 323.6 |

**Table S3. Parameters considered in the vampire and cavity pinning feature design in STOMP.**

| Parameter | Convex "Vampire" Pinning Feature | Concave "Cavity" Pinning Feature |
|---|---|---|
| Width between pins, $w_1$, $w_2$ (mm) | 0.8 | 1.2 |
| Vampire angle, $\alpha$ (degrees) | 5 to 80 | – |
| Cavity angle, $\beta$ (degrees) | – | 25 to 130 |
| Channel height, $h$ (mm) | 3.5 | |
| Collagen density (kg/m$^3$) | 958 | |
| Surface tension (mN/m) | 72 | |
| Average static contact angle of collagen on 3D printed resin, $\theta$ (degrees) | 30 ± 3.5 | |
| Lowest measured contact angle (degrees) | 15 | |
| Highest measured contact angle (degrees) | 48 | |
| Gravitation constant (m/s$^2$) | 10 | |

**Movie S1 (separate file).** Patterning three regions with STOMP. Purple-colored agarose is pipetted first in the two outer regions followed by yellow-colored agarose in the middle region. This video was used to generate Fig. 1g.

**Movie S2 (separate file).** Successful pinning of 23 μL of a 5 mg/mL precursor collagen solution in convex "vampire" pinning feature with $\alpha = 20°$. Prior to testing, the STOMP device was treated with 1% BSA. Pinning was considered successful if the fluid stayed pinned for 10 s after pipetting. This video was used to generate Fig. 1e.

**Movie S3 (separate file).** Loss of pinning of 23 μL of a 5 mg/mL precursor collagen solution in convex "vampire" pinning feature with $\alpha = 45°$. Prior to testing, the STOMP device was treated with 1% BSA. This video was used to generate Fig. 1e.

**Movie S4 (separate file).** Successful pinning of 23 μL of a 5 mg/mL precursor collagen solution in concave "cavity" pinning feature with $\beta = 40°$. Prior to testing, the STOMP device was treated with 1% BSA. Pinning was considered successful if the fluid stayed pinned for 10 s after pipetting. This video was used to generate Fig. 1f.

**Movie S5 (separate file).** Loss of pinning of 23 μL of a 5 mg/mL precursor collagen solution in concave "cavity" pinning feature with $\beta = 100°$. Prior to testing, the STOMP device was treated with 1% BSA. This video was used to generate Fig. 1f.

**Movie S6 (separate file).** Spontaneous beating of unpaced control engineered heart tissue (EHT) patterned with STOMP in GFP channel. This EHT contains cardiomyocytes, HS27a stromal cells, and HS5-GFP cells in the two outer regions and cardiomyocytes, HS27a stromal cells, and HS5-mCherry cells in the middle region. This video was used to generate Fig. 3b.

**Movie S7 (separate file).** Spontaneous beating of unpaced control engineered heart tissue (EHT) patterned with STOMP in mCherry channel. This EHT contains cardiomyocytes, HS27a stromal cells, and HS5-GFP cells in the two outer regions and cardiomyocytes, HS27a stromal cells, and HS5-mCherry cells in the middle region. This video was used to generate Fig. 3b.

**Movie S8 (separate file).** Patterning degradable channel wall with tissue region. Pink-colored agarose is pipetted first into the degradable channel wall region within the STOMP device followed by green-colored agarose in the main tissue region of the STOMP device. This video was used to generate Fig. 5e.

**Movie S9 (separate file).** Removing STOMP device containing 3T3 cells laden in poly(ethylene glycol) (PEG) after degrading channel walls.

**Design File S1 (separate file).** 3.5 mm height 1.2 mm width 3 region convex “vampire” pinning feature STOMP ($\alpha$=60°, used in Fig. 1, Fig. 3, and generating EHTs).

**Design File S2 (separate file).** 3.5 mm height 1.2 mm width 3 region convex “vampire” pinning feature STOMP ($\alpha$=10°, used in Fig. 2)

**Design File S3 (separate file).** 3.5 mm height 1.2 mm width 3 region convex “vampire” pinning feature STOMP ($\alpha$=20°, used in Fig. 2)

**Design File S4 (separate file).** 3.5 mm height 1.2 mm width 3 region convex “vampire” pinning feature STOMP ($\alpha$=30°, used in Fig. 2)

**Design File S5 (separate file).** 3.5 mm height 1.2 mm width 3 region convex “vampire” pinning feature STOMP ($\alpha$=45°, used in Fig. 2)

**Design File S6 (separate file).** 3.5 mm height 1.2 mm width 3 region concave “cavity” pinning feature STOMP ($\beta$=40°, used in Fig. 2)

**Design File S7 (separate file).** 3.5 mm height 1.2 mm width 3 region concave “cavity” pinning feature STOMP ($\beta$=60°, used in Fig. 2)

**Design File S8 (separate file).** 3.5 mm height 1.2 mm width 3 region concave “cavity” pinning feature STOMP ($\beta$=90°, used in Fig. 2)

**Design File S9 (separate file).** 3.5 mm height 1.2 mm width 3 region concave “cavity” pinning feature STOMP ($\beta$=100°, used in Fig. 2)

**Design File S10 (separate file).** 3.5 mm height 1.2 mm width 3 region concave “cavity” pinning feature STOMP ($\beta$=120°, used in Fig. 2)

**Design File S11 (separate file).** 2 mm height 1 mm width 3 region concave “cavity” pinning feature STOMP ($\beta$=90°, used in Fig. 4 and generating PTCs)

**Design File S12 (separate file).** 2 mm height 1 mm width single region STOMP (inner core region device, used in Fig. 5)

**Design File S13 (separate file).** 2 mm height 1 mm width 2 region convex “vampire” pinning feature STOMP ($\alpha$=20°, used in Fig. 5)

**Design File S14 (separate file).** 2 mm height 1 mm width 3 region convex “vampire” pinning feature STOMP ($\alpha$=20°, used in Fig. 5)

**Design File S15 (separate file).** 4 mm height 1.9 mm width single region STOMP (outer region device, used in Fig. 5)

**Design File S16 (separate file).** Degradable v-shaped channel walls STOMP device (used in Fig. 5)

***All design files will be available in a public repository.**
